# Using artificial intelligence to optimize ecological restoration for climate and biodiversity

**DOI:** 10.1101/2025.01.31.635975

**Authors:** Daniele Silvestro, Stefano Goria, E-Ping Rau, Renato Augusto Ferreira de Lima, Ben Groom, Piotr Jacobsson, Thomas Sterner, Alexandre Antonelli

## Abstract

The restoration of degraded ecosystems is critical for mitigating climate change and reversing biodiversity loss. Depending on the primary objective – such as maximizing carbon sequestration or protecting threatened species – and within the boundaries of budget limitations, different spatial priorities have been identified at global and regional scales. Funding mechanisms to support such work comprise public sources, philanthropy, and the private sector, including the sale of carbon and biodiversity credits. However, effectively exploring trade-offs among restoration objectives and estimating the price of biodiversity and carbon credits to design financially viable projects remain scientifically challenging. Here we harness the power of artificial intelligence implemented in the software CAPTAIN to identify spatial priorities for ecological restoration that maximize multiple objectives at once and to allow a robust evaluation of biodiversity and climate outcomes. We find through a series of simulations that even low-to-moderate consideration of biodiversity in the design of restoration projects leads to the selection of restored areas that disproportionately improve the conservation of threatened species, while the carbon captured is only slightly reduced. We propose the bundling of biodiversity and carbon credits to support restoration efforts even in areas previously excluded for economic reasons. Applying our methodology to plant diversity in the Atlantic Forest of eastern South America, one of the most biodiverse and threatened ecosystems globally, we demonstrate its potential to guide restoration and to quantify trade-offs between multiple optimization scenarios. Our study provides a robust, scalable method to maximize the joint benefits of ecological restoration for climate and nature.

## Introduction

Ecological restoration is one of the most powerful nature-based solutions to tackle the twin planetary crises of biodiversity loss and climate change^1^. In recognition of its importance, the United Nations has declared 2021–2030 the Decade on Ecosystem Restoration and it motivates the commitment under Target 2 of the Kunming-Montreal Global Biodiversity Framework – adopted by 196 countries – that “*by 2030 at least 30 per cent of areas of degraded terrestrial, inland water, and marine and coastal ecosystems are under effective restoration*”. In addition to protecting natural ecosystems - the preferred option whenever possible - well-designed and implemented ecological restoration can capture and store carbon, increase populations of threatened species and reinstate or strengthen many of nature’s contributions to people, helping us achieve several of the Sustainable Development Goals^2–4^.

Despite the multiple benefits of restoration and its urgency in the face of global environmental change, two major challenges hamper its implementation. Firstly, much like prioritizing areas for protection^5^, there are clear trade-offs when designing restoration projects aiming to maximize carbon capture or biodiversity recovery^6^. For example, the planting of exotic trees across Africa, although they grow quickly and capture large amounts of carbon (in principle helping to mitigate climate change, if the biomass produced is retained long-term), has led to detrimental effects on native biodiversity^7^. Trade-off effects have been widely discussed in the literature e.g.^8,9^, but to our knowledge no tools have yet been developed to allow their quantification while taking into account the complexity of the real world – including the spatial heterogeneity in biodiversity, carbon storage potential, costs, and their changes over time.

Secondly, there is a major funding gap – estimated to exceed 600 billion USD annually^10^ – to implement the Global Biodiversity Framework, including its restoration target. Given that public funding from institutions such as the Global Environment Facility, which provides funding of roughly USD 1.3 billion per year, falls far short of reaching the required levels, many restoration projects today are supported instead by the sales of carbon credits, which can be voluntary or mandatory depending on the country. Alas, the simplicity of carbon credits, where one credit equals one tonne of atmospheric carbon captured, regardless of location, is also a disadvantage from the viewpoint of biodiversity. As market forces push trade towards low prices, they incentivize restoration projects that minimize costs (such as reforestation on marginal and highly degraded land) and focus exclusively on maximum carbon recovery, even at the cost of biodiversity benefits. To improve this situation, carbon credits are being designed to include additional benefits such as biodiversity, and in parallel, biodiversity credits are emerging following strict principles for high integrity and governance^11,12^. Quantifying the cost relationship between carbon and biodiversity credits to make them financially interchangeable (allowing, for instance, a restoration project to become viable by selling biodiversity credits rather than carbon credits) can create economic opportunities for bundling carbon and biodiversity credits for the same site, while avoiding double claims. This strategy can enable the restoration of ecosystems in expensive areas, which have so far been excluded due to insufficient profit margins^13^.

Here, we propose an approach based on multi-objective reinforcement learning to address these shortcomings. Specifically, we: i) show through simulations that this approach can help spatially guide restoration programs while monitoring their outcomes over time; ii) explore trade-offs and synergies that exist between restoration efforts focused on carbon, biodiversity and costs, through multi-objective optimization to identify socially optimal solutions; iii) link our predictions to the emerging concept of biodiversity credits alongside carbon credits to help design financially viable restoration programs; and iv) run our optimization framework as a proof-of-concept on a large empirical dataset from the Atlantic Forest in South America. Our results show that identifying trade-offs between carbon and biodiversity is crucial to evaluate the potential impact of biodiversity credits and to achieve restoration programs that maximize benefits for both climate and biodiversity.

## Results

### A reinforcement learning framework for restoration

We developed an approach to optimize restoration planning building upon the reinforcement learning software CAPTAIN (Conservation Area Prioritization Through Artificial INtel- ligence)^14^. The implementation is based on a spatially explicit dynamic *environment* characterized by climatic features, species and their geographic ranges, anthropogenic disturbances, and restoration costs. The disturbance could be any kind of anthropogenic activity including settlements, roads, farming or logging on land and fishing or mining in the sea, while costs represent direct restoration costs and opportunity costs, e.g. if a restored area can no longer be used for farming.

The other element of the CAPTAIN framework is an *agent* (i.e. a virtual representation of a manager or social planner) who monitors the environment and implements a restoration program pursuing one or multiple pre-defined restoration targets. The action performed by the agent is the selection of areas within which the disturbance is reduced and natural restoration can occur. The agent uses a neural network to map the current state of the environment to a selection of areas for restoration. The environment evolves over time through a number of steps (e.g. years, here set to 30) during which multiple processes take place, including dispersal and mortality among species, changes to climate or disturbances, and the establishment of restoration areas selected by the agent. The intervention of the agent is assigned a reward based on its outcome, for instance its contribution to preventing species extinctions, and models are optimized to maximize the reward based on the outcomes of the agent’s actions.

We implemented a multi-objective reinforcement learning algorithm to evaluate the ability of this approach to optimize restoration toward multiple goals, specifically targeting the net carbon sequestration and reduction in species extinction risks, while incorporating restoration costs (see *Methods*, Equations 6–12). Here, we measured extinction risks in a 5-tiered classification inspired by the International Union for Conservation of Nature (IUCN)^15^ and ranging from Least Concern to Critically Endangered. The model is highly flexible and can be tuned to optimize other parameters, such as biodiversity metrics based on richness, abundance and function. We optimized five models to get different spatial prioritizations, which we hereafter refer to as *policies*^16^, using differently weighted rewards. Specifically: Policy 1) focused on carbon, Policy 2) included extinction risk with a small weight, Policy 3) weighted carbon and biodiversity equally, Policy 4) assigned a smaller weight to carbon, and Policy 5) focused on biodiversity (Table 2). All policies shared the same weight on the budget, rewarding the reduction of restoration costs. All policies were constrained to restore the same total area, which makes the biodiversity and carbon outcomes more comparable among policies, but which means that they may differ in implementation costs. We then compared the predicted restoration outcomes of different policies, in terms of selected areas, resulting net carbon sequestration, and impact on species extinction risks. This allowed us to evaluate a gradient of restoration policies ranging from climate- to biodiversity-optimal. In all plots we use a color gradient from brown to green to indicate policies with rewards shifting from carbon to biodiversity.

### Optimization of restoration policies and trade-offs

Our simulations of biodiversity used to optimize restoration policies and predict their outcomes revealed that the five policies resulted in substantial spatial differences in the prioritization of restoration areas and led to widely different carbon and biodiversity outcomes. While model training and simulations were based on the same arbitrarily chosen geographic region (Figs. S2-S3), all datasets differed in the distribution of simulated species, threats, and costs, in order to cover a wide range of possible real-world scenarios. Consequently, the placement of the selected restored areas varied among them. However, at the level of each simulation, the policies clearly prioritized different areas (Fig. S1).

Policy 1 favored the restoration of areas that resulted in the highest net carbon gain (24% on average after 30 time steps; Fig. 1a). In contrast, Policy 5 prioritized areas that contributed the most to reducing species extinction risk, therefore considering threats, species richness, and complementarity of species composition (Fig. S1). The result was a more modest carbon gain (just above 11% on average; Fig. 1a) but a substantially stronger reduction of the extinction risk compared with Policy 1. For instance, the average number of Critically Endangered species at the end of the simulation was 36% lower under Policy 5 compared to Policy 1 (Fig. 1b). Similarly, the number of species classified as Endangered decreased on average by 73% when giving priority to biodiversity over carbon. These experiments also showed that while carbon accumulated through time following near-linear trajectories, extinction risks followed more complex paths (Fig. 1b-f), reflecting the fact that species moved across risk categories and that their conservation status was determined by their ability to re-colonize restored areas based on their dispersal ability, their sensitivity to disturbance, and connectivity among suitable areas.

**Figure 1:**
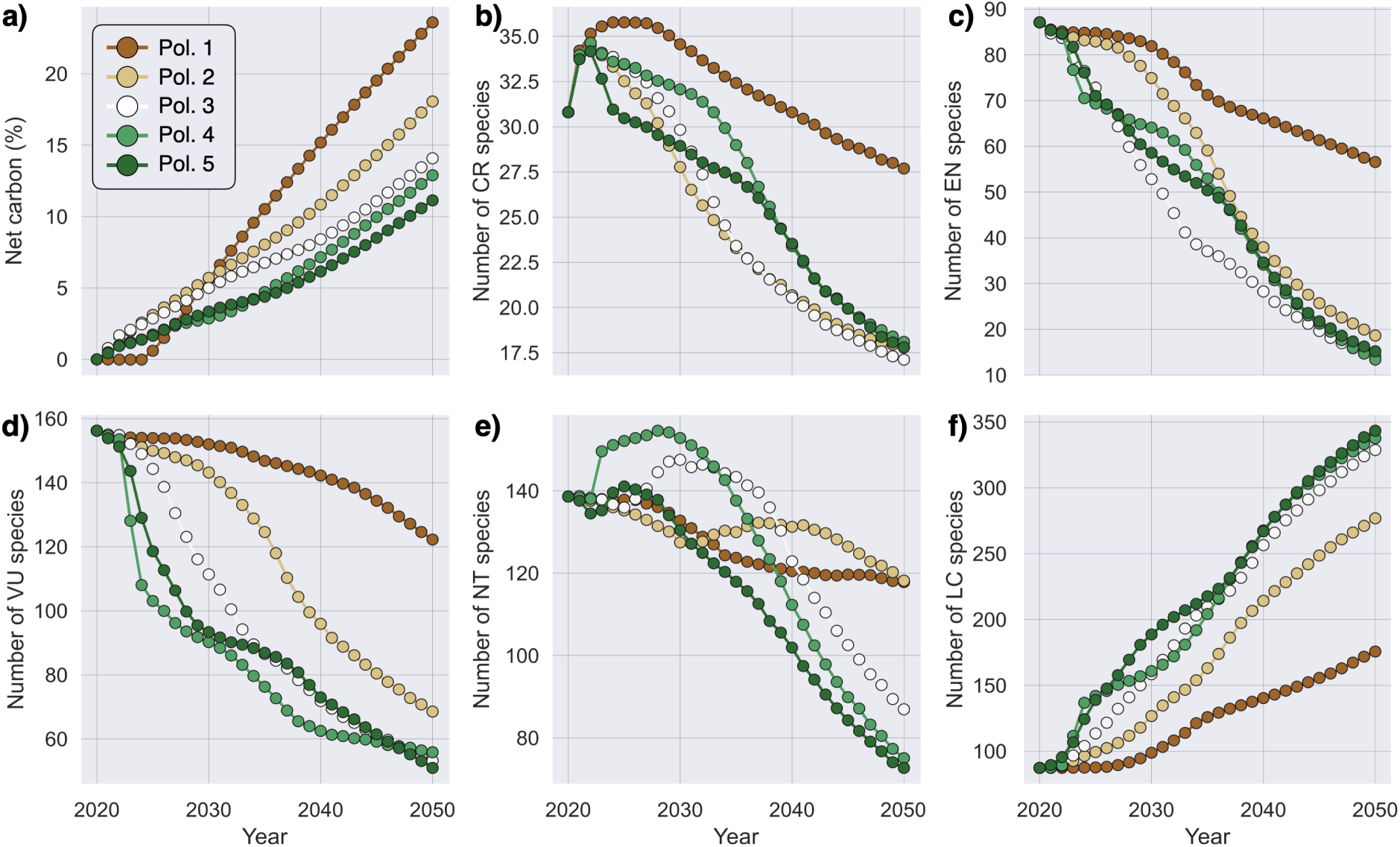
Accumulation of carbon and evolution of extinction risks across 30 time-steps averaged over the 100 simulations. The results are shown for Policies 1–5, which differ in the relative weight assigned to carbon sequestration (highest in Policy 1) and reduction of extinction risks (highest in Policy 5). The plots show that when biodiversity is factored in, the carbon accumulation curve reaches lower levels decreasing from 24% to 11% (a). This trend is counterbalanced by a substantial improvement of species conservation status, with a reduction of Critically Endangered and Endangered species of 36 and 73%, respectively, from Policy 1 to 5 (b, c). Meanwhile, the number of Least Concern species increased by 51% (f), highlighting the substantial impact of biodiversity-targeted restoration in our simulations. The time labels used here are arbitrary.

The distributions of outcomes at the end of the 30 time-step restoration show a substantial level of consistency across 100 simulations (Fig. S6). The net carbon gain in Policy 1, which was trained without considering a biodiversity reward, resulted in an average 2.2-fold increase (standard deviation among simulations: ±0.26) in carbon storage compared to Policy 5, which did not include carbon in the reward (Fig. 2a). Policies with mixed rewards revealed a gradient of outcomes, with net carbon gain gradually decreasing when setting the relative carbon reward to lower values (Figs. 1b, 2a). In contrast, the biodiversity outcomes revealed drastic improvements as soon as a non-zero weight was assigned to the biodiversity reward, with Policies 2–5 all resulting in similar numbers of species with reduced extinction risk (Fig. 2b), while Policy 1 resulted in significantly worse impacts on extinction risks compared to most other policies (Figs. 2, S6b). For instance, while on average 30% of the species decreased their extinction risk under Policy 1 (150 out of the 500 simulated species), the percentage rose to 71–76% (354–378 species) under all other policies (Fig. S6b). The fraction of Critically Endangered species remained virtually unchanged under Policy 1, while it decreased on average by ∼29% under Policy 5, and by 27% under all other policies (Fig. S6d). When combining all threatened species (CR, EN, VU), these decreased by about 25% under Policy 1 (i.e. ∼68 species) but dropped by 62–69% (i.e. 170–190 species) under the others (Fig. S6i), thus showing a 2.6-fold improvement. These results show that even when biodiversity outcomes are given a smaller weight compared to carbon in the optimization of the restoration policy (e.g., Policy 2), the impact on extinction risk reduction is large. Finally, the costs of implementation of the different policies, i.e. the cumulative cost of all restored cells, were relatively similar, with changes between 4 and 17% among policies (Fig. 2c). The variation results from setting a constraint on the number of restored cells rather than on the total budget, a choice that makes the biodiversity and carbon outcomes more comparable among policies. Yet, the low variation in total costs reflects the fact that the same weight was assigned to the budget reward in the optimization of all policies (see *Methods*).

**Figure 2:**
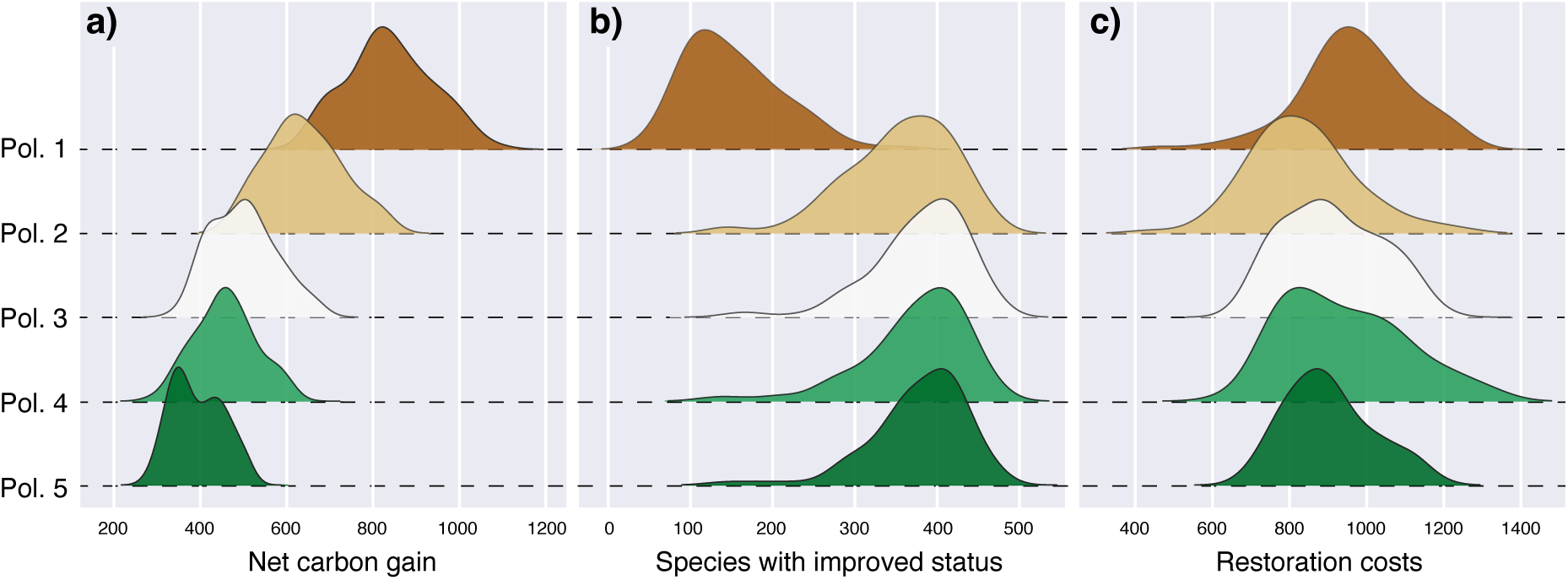
Summary statistics of key outcomes from 100 simulations run under five trained restoration models, Policies 1–5, ranging in their objective from carbon-only (1), through mixed objectives (2–4), to biodiversity-only (5). The net carbon (a) represents the net gain in amount of carbon (here expressed in arbitrary units – see *Methods*) in the system after 30 time-steps during which 3,000 cells were restored. As the weight assigned to the carbon reward was lowered from 0.5 to 0 in favor of the biodiversity reward, the net carbon gain gradually decreased, although all restoration policies resulted in a net gain. In contrast, the number of species with improved conservation status, i.e. lower extinction risk, drastically increased as soon as a non-zero weight was assigned to the biodiversity reward (Policies 2–5; b). The implementation cost (here expressed in arbitrary units – see *Methods*) was similar across all policies (c), reflecting the fact that all models were trained with the same budget reward.

Additional experiments showed that our results are robust to changes in the architecture of the neural network underlying the different policies (Table S3). In contrast, analyses of the simulated data under simple rule-based selection of the cells to be restored (e.g. prioritizing cells with the highest ratio between number of threatened species and restoration cost) yielded substantially worse outcomes (Table S3; *Supplementary Results*). This indicates that the neural network is better able to capture the dynamics of the environment to select restoration areas.

### Carbon credits and the impact of biodiversity credit valuations

Our analyses revealed that different restoration policies lead to vastly divergent carbon and biodiversity outcomes while maintaining comparable implementation costs. To evaluate their financial feasibility, we first performed a break-even analysis assuming a carbon-only credit market. We set the carbon credit value (*χ*) such that the cumulative revenue from Policy 1, which maximizes carbon sequestration, exactly matched its total restoration costs (Eq. 14; Fig. 3a). Under this baseline, all other policies incurred net financial losses, ranging from -10% for Policy 2 to approximately -50% for Policy 5, as they prioritize biodiversity at the expense of carbon sequestration.

**Figure 3:**
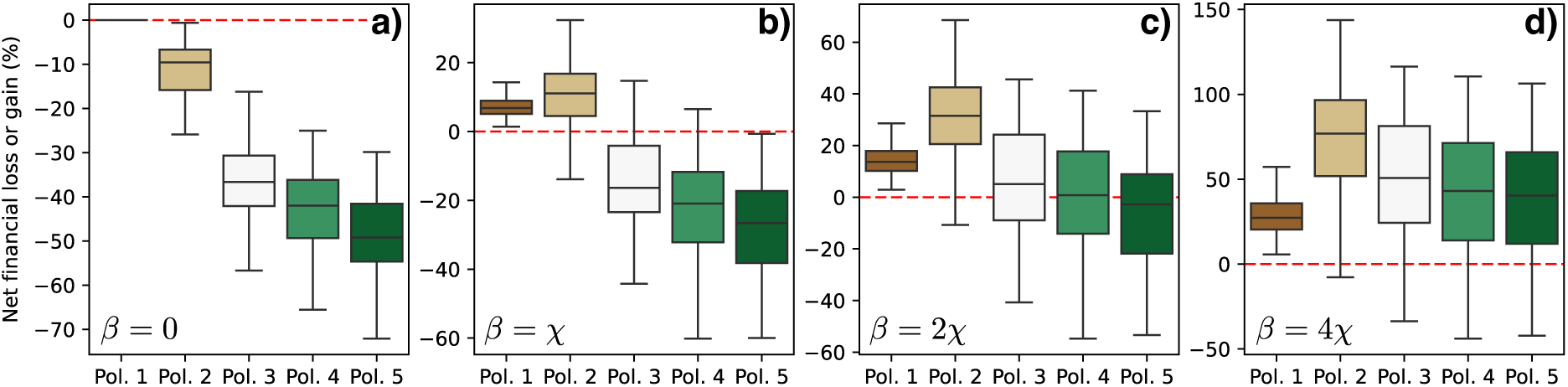
Financial outcomes of restoration policies (net percentage loss or gain, i.e. Φ in Equation 15) under different valuations of biodiversity credits. Negative values indicate a net economic loss where restoration costs exceed the cumulative value of carbon and biodiversity credits, while positive values indicate a net gain. The results are shown when only carbon credits (*χ*) are included (i.e. *β* = 0; a), which we used as a baseline of net 0 financial outcome to show how biodiversity credits could contribute value to the simulated restoration programs. The results in b–d) show that when biodiversity credits are valued at *β* = *χ,* 2*χ* and 4*χ*, respectively, all policies become financially viable (Φ *>* 0).

We then explored the financial impact of bundling carbon and biodiversity credits. Recogniz- ing that there is currently no global consensus on the unit of a biodiversity credit^17,18^, we adopted a performance-based metric focused on tangible ecological outcomes. For this demonstration, one biodiversity credit (*β*) corresponds to one species transitioning from a threatened category to a non-threatened one by the end of the simulation. Without implying that this should be preferred over alternative definitions, we used this measure to evaluate the potential effects of biodiversity credits on the financial viability of different restoration strategies.

Our experiments with varying relative valuations (*β* = 1*χ* to 4*χ*) demonstrated that biodiversity credits can effectively level the financial barriers to multi-objective restoration. When the value of one biodiversity credit was set equal to one carbon credit (*β* = *χ*), Policy 2 -which maintains a carbon-oriented focus while integrating a moderate biodiversity weight- became profitable, yielding gains above 10%. Further increasing the relative value of biodiversity (*β* = 2*χ* to 4*χ*) improved the financial outcomes for all policies (Fig. 3c, d). Notably, Policy 2 consistently outperformed the others, reflecting the non-linear biodiversity gains achievable when even moderate weight is given to conservation (Fig. S6b).

Finally, we derived the “equivalence value” of a biodiversity credit, i.e. the price required for a biodiversity-aware policy (Policies 2–5) to match the net profit of the carbon-optimal Policy 1 (Eq. 17). We found that Policies 3–5 reached parity with Policy 1 at valuations of *β* = 2.6 − 3.5*χ*, whereas for Policy 2, a value of *β* = 0.8*χ* was sufficient (Table S1). This implies that if a biodiversity credit is valued at just 1.2 times a carbon credit, Policies 1 and 2 become economically indistinguishable.

While we acknowledge these valuations are currently expressed in arbitrary units, our framework provides a reproducible path for deriving the real-world value of biodiversity credits based on defined ecological targets. In contrast, assigning a null value to biodiversity (*β* = 0) leaves limited options to target biodiversity in restoration programs without drastically reducing the financial viability of the policy.

### Restoration priorities and trade-offs in the Atlantic Forest

We tested our multi-objective reinforcement learning model on an empirical dataset from the Atlantic Forest – one of the world’s most biodiverse and threatened ecosystems^19,20^. Our analysis was based on a recently published dataset^19^, which included *>*3,800 species of trees and shrubs. The data included species distribution models based on approximately 900,000 curated occurrences, an approximation of species-specific relative contribution to above-ground carbon, and extinction risks based on the IUCN Red List classification scheme. We also incorporated a proxy for current disturbance levels based on land use, and restoration costs based on accessibility, elevation and value of agricultural production (see *Methods and Supplementary Information*; Fig. S12).

We ran three models based on Policies 1 (carbon-focused), 3 (equal weights for carbon and biodiversity), and 5 (biodiversity-focused), as described above. Simulation experiments carried out outside of the region used for model training showed that the optimized policies can be applied to different contexts with reasonable performance (Fig. S1, S7; *Supplementary Methods and Results*). However, they also suggest that models achieve higher and more consistent performance when applied within the region used in their optimization. For this reason, we fine-tuned the simulation-trained models with the Atlantic Forest data, to identify the 30% with highest restoration priority (i.e. 4,800 of the 16,263 spatial units considered here). The training was based on weighted rewards linked to carbon, species protection, and costs computed as defined in Equations 6–12. As some of the parameters of analysis were not available or incomplete (such as species-specific growth rates, sensitivity to disturbance, and extinction risks), we performed 30 runs to account for variation in the imputation of missing data (see *Methods*).

The selected areas showed substantial differences among policies, highlighting partial mismatches between predicted restoration potential in terms of carbon and biodiversity outcomes (Fig. 4a). Specifically, 63% of the areas selected under a carbon-focused policy overlapped with those selected under a policy that accounted for both carbon and biodiversity, while the overlap dropped to 34.5% when comparing against a policy focused only on biodiversity outcomes. Meanwhile, Policies 3 and 5 overlapped by 53.3%. The estimates obtained across different runs showed that variation in the imputation of missing data has a non-negligible impact on the results, although selected spatial priorities were generally consistent among replicates. The overlap across runs was 94% for Policy 1, 80% for Policy 3, and 74% for Policy 5, suggesting that missing data had a greater impact on restoration priorities targeting extinction risk compared to carbon (Fig. S9).

**Figure 4:**
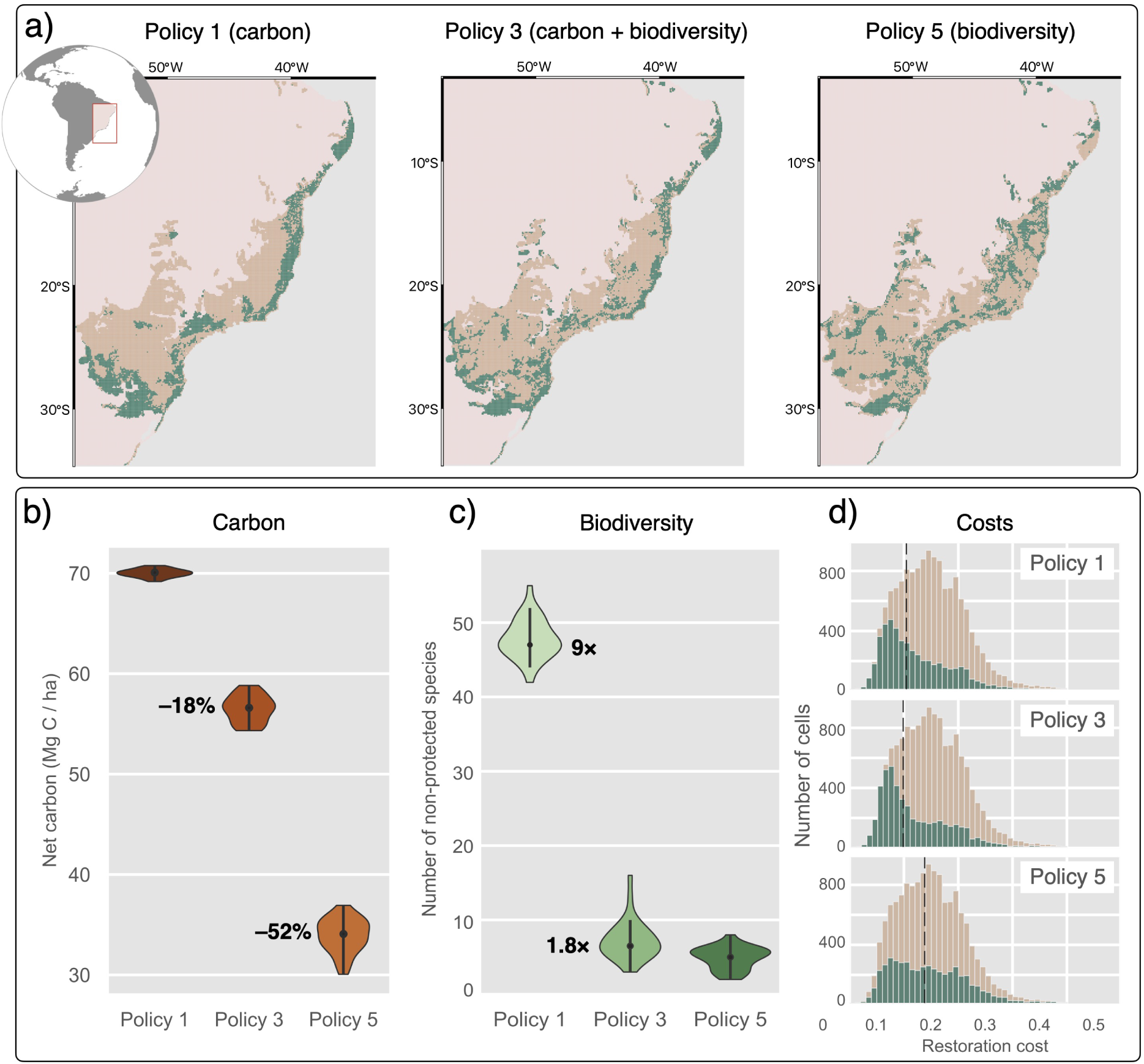
Results of CAPTAIN optimizations in the Atlantic Forest based on a dataset including habitat suitability maps for 3,789 species of trees and shrubs, and proxies for costs and land use. The extent of the considered area is shown in brown, while the cells selected for restoration are shown in green. The data are analyzed under three policies (as defined in Table 2) that were focused on carbon (Policy 1), reduction of extinction risks (Policy 5) or both (Policy 3). All models also used a budget reward for costs and were constrained to restore the same number of grid cells (i.e. 30% of the total). The selected areas, after fine-tuning the simulation-trained models with the empirical data, are only partly overlapping, indicating substantial trade-offs among priorities (a). The predicted outcomes also vary substantially among policies in terms of expected amount of net carbon sequestration (approximated as described in the *Methods*) (b) and number of species occurring only outside of protected areas (c). The violin plots show the outcomes of 30 runs that incorporate uncertainties in the imputation of missing data (see *Methods* and Fig. S9a–c). All policies selected cells with a wide range of costs (d).

The outcomes of our inference were evaluated through comparisons between the predicted changes in carbon and species extinction risks obtained through restoration and the starting condition (baseline) of current disturbance and threat levels. This comparison is essentially equivalent to a business-as-usual counterfactual because the ecosystem is simulated to be at equilibrium with no changes in disturbance, extinction risks, or species abundances through time, in the absence of restoration actions. The results reflected the objectives the models had been optimized for, showing clear trade-offs between the potential for carbon sequestration through restoration and the ability to protect species and reduce extinction risks. Using species-specific proxies for above-ground carbon and calibrating the average carbon per spatial unit to 98.5 tonnes per hectare (Mg C / ha) in the Atlantic Forest^21^, we found that Policy 1 led to an average predicted net carbon sequestration (approximated as described in the *Methods*) ranging from 69.3 to 70 Mg C / ha, across 30 runs (Fig. 4b). For comparison, Policies 3 and 5 led to a predicted carbon sequestration of 54.3–58.8 and 30.1–36.9 Mg C / ha, respectively, thus leading to a decrease in the order of 20 to 50%.

In contrast, areas selected under Policy 1 left 42–55 species under no protection, a number that dropped to 3–16 under Policy 3 and to 2–8 under Policy 5 (Fig. 4c). Under Policy 1, between 15 and 38 of the 420 Critically Endangered and Endangered species included in the dataset (391–453 across the 30 runs) remained in these high-risk categories at the end of the simulation. This indicates that the selected areas failed to restore a sufficient fraction of species’ suitable habitats (set to 25% for CR species and 10% for EN species) required to move to a lower risk category under our thresholds (Table 1). The median number of high risk species dropped to 0 under Policy 3, albeit with some degree of variability among runs (0–4), which reduced further under Policy 5, where at most 1 species was left in that group.

**Table 1:**
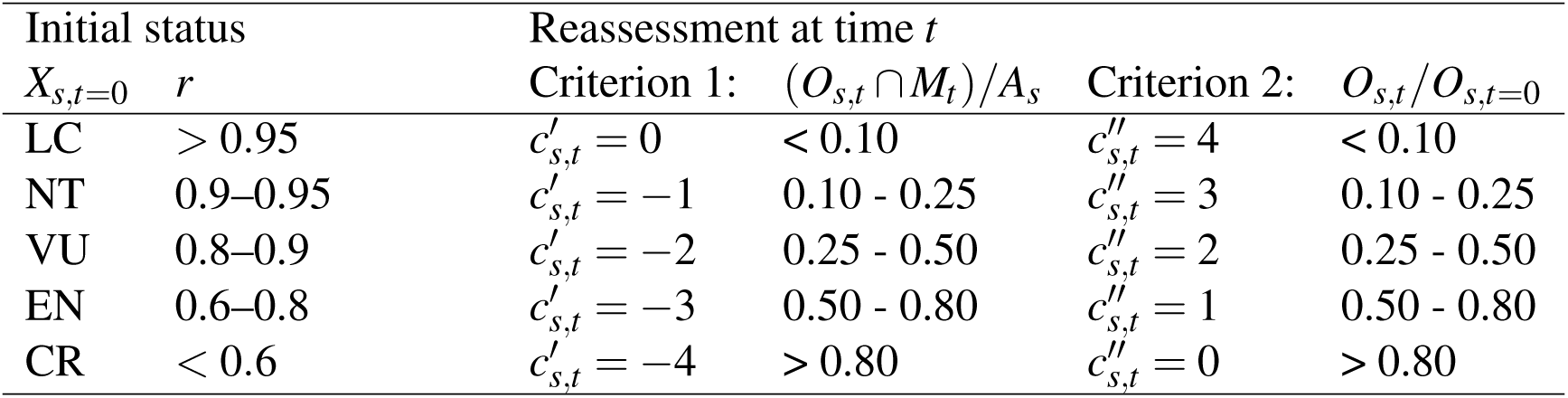
Thresholds used to set the initial species extinction risks (*X_s,t_*_=0_, only applied in simulated data and replaced with Red List assessment when using empirical data) and their reassessment at time *t* based on their level of protection (Criterion 1) and population trends (Criterion 2). The re-assessed extinction risk is 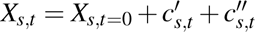

| Initial status | | Reassessment at time $t$ | | | |
| --- | --- | --- | --- | --- | --- |
| $X_{s,t=0}$ | $r$ | Criterion 1: $(O_{s,t} \cap M_t)/A_s$ | | Criterion 2: $O_{s,t}/O_{s,t=0}$ | |
| LC | $> 0.95$ | $c'_{s,t} = 0$ | $< 0.10$ | $c''_{s,t} = 4$ | $< 0.10$ |
| NT | $0.9-0.95$ | $c'_{s,t} = -1$ | $0.10 - 0.25$ | $c''_{s,t} = 3$ | $0.10 - 0.25$ |
| VU | $0.8-0.9$ | $c'_{s,t} = -2$ | $0.25 - 0.50$ | $c''_{s,t} = 2$ | $0.25 - 0.50$ |
| EN | $0.6-0.8$ | $c'_{s,t} = -3$ | $0.50 - 0.80$ | $c''_{s,t} = 1$ | $0.50 - 0.80$ |
| CR | $< 0.6$ | $c'_{s,t} = -4$ | $> 0.80$ | $c''_{s,t} = 0$ | $> 0.80$ |

The total cost of Policy 3 was about 1% lower than that of Policy 1 and 16% lower than Policy 5. However, all policies showed that the cost of selected cells covers most of the range of affordable cells (that is, excluding urban areas; Fig. 4d). This indicates that the restoration areas optimized by CAPTAIN still included expensive cells despite a negative reward associated with implementation costs.

We carried out sensitivity tests to assess the impact of potential error in carbon estimates and opportunity costs (see *Supplementary Methods and Results*). For these analyses we simulated random error in the carbon and cost data and repeated the analyses across 30 runs. The results showed that the selected priority areas remained broadly consistent with those obtained with the original dataset (Figs. 4a, S9). However, the simulated error led to a lower average overlap across the priority areas selected in different runs, which decreased to 51–94% (from the 74–94% variation in analyses without error; see *Supplementary Methods and Results*). These results show that while some of the selected areas are robust to error in the underlying data, others are more sensitive. This demonstrates the importance of obtaining high quality data to reduce uncertainties in real-world applications of CAPTAIN in restoration planning. The analyses show that our approach can be used to explicitly tackle, rather than ignore, deficiencies and uncertainties inherent in real-world data, in order to quantify the robustness of the results. Meanwhile, regions where error leads to shifting prioritization highlight areas that might require additional verification of the cost, carbon, and biodiversity data, to achieve a robust optimization of restoration planning.

## Discussion

### Reinforcement learning as a tool for guiding restoration

Recent work has shown that reinforcement learning holds potential in the context of dynamic ecological settings, for instance in guiding patrolling of protected areas^22^, selecting conservation areas^14,23^, improving their connectivity^24^, or modeling ecosystem dynamics^25^. Here, we demonstrated the power of multi-objective reinforcement learning to optimize ecological restoration strategies targeting climate- and biodiversity-positive outcomes while accounting for costs of implementation. Using weighted rewards in the training phase allowed us to calibrate models that explore the boundary of optimality^26^ between net carbon gain and reduction in extinction risks obtained through restoration, while keeping the extent of restored area constant. This simulation and optimization framework allows us to evaluate the synergies and trade-offs among restoration policies with different objectives, as also shown in applications in conservation planning^27,28^.

As our models are trained within a biologically and environmentally dynamic environment, the solutions incorporate the effects of connectivity among restored areas, which favor migration processes and range expansion underlying the recovery of our simulated species. They can also account for species-specific features such as climatic niche, potential for carbon sequestration, growth and dispersal rates, and sensitivity to anthropogenic disturbance. The simulations show that the spatial distribution of biodiversity and carbon priorities differs, as observed in real-world scenarios^5,29^. This mismatch means that a restoration program focusing on the maximization of carbon will likely not result in the best biodiversity outcome, and vice versa.

Our analysis of empirical data from the Brazilian Atlantic Forest confirms the trends observed with simulated data. A restoration program optimized exclusively for carbon sequestration fails to adequately protect biodiversity, even when based on natural restoration as opposed to monoculture plantations. We have not considered monocultures in our study, given their well-known harmful impact on the environment (e.g.,^7^) and since ecological restoration projects in Brazil do not use single-species plantations. On the other hand, including the reduction of extinction risks into the optimization will typically also lead to a reduction in net carbon gain. This is because threatened species are not necessarily the fastest-growing ones (such as the giant timber trees in the genus *Cariniana*, which take decades to reach maturity) or those with largest biomass (such as the many treelets in the genus *Myrcia*).

While no solution can thus achieve maximum levels of biodiversity outcomes and carbon sequestration simultaneously, our empirical analysis for the Atlantic Forest shows that the loss of biodiversity protection inflicted by focusing only on carbon would be substantially larger (ca. 7 times more species under no protection) than the reduction of carbon sequestration (ca. 18% decrease) due to the incorporation of biodiversity outcomes (see Policies 1 vs 3 in Fig. 4b,c). The results also indicate that the incorporation of carbon outcomes leads to only a minor loss of biodiversity benefits (Policies 3 vs 5 in Fig. 4b,c). Although our implementation gives substantial flexibility in creating realistic models of the environment’s dynamics, ultimately the application of this methodology to real-world restoration planning depends on the accuracy of the data fed into the model. This will require moving beyond the approximations for cost, land use, and habitat degradation used in our proof-of-concept implementation and the integration of additional constraints, for instance linked with property boundaries and restoration feasibility.

Within the limits of available data, our multi-objective reinforcement learning framework provides a robust tool to explore trade-offs between the arguably most important objectives of ecological restoration: carbon sequestration and biodiversity uplift, allowing for the optimization of these potentially conflicting metrics. However, we acknowledge that final decisions of where restoration efforts will occur depend on additional considerations other than the ones explored here - such as the local interests and support of Indigenous and local communities, political commitments at national and regional levels, availability of seedling supply, accessibility, human capacity, and other resources.

### The potential of bundled carbon and biodiversity credits

Our simulation experiments show that an evaluation of restoration outcomes and their associated costs can be used to forecast the impact of biodiversity credits on program feasibility. We show that the financial sustainability of restoration policies targeting both carbon sequestration and extinction risk is highly sensitive to the valuation of biodiversity relative to carbon. Crucially, our results indicate that even a modest premium assigned to biodiversity outcomes can create incentives for multi-objective restoration that addresses both carbon sequestration and extinction risks.

In principle, this suggests that high-integrity carbon credits, which explicitly consider and demonstrate co-benefits such as biodiversity gain, could help bridge the funding gap for projects that effectively achieve both climate and ecological goals. These carbon credits could be traded at a premium price to compensate for the costs of activities that protect, restore and monitor biodiversity, as well as to generate additional benefits that could be shared within the community involved in the conservation projects. In practice, ecological restoration companies frequently report that such added costs remain too high to be competitive in the global carbon market, pushing restoration projects to marginal, inexpensive land, where ecological potential may be lower. This underscores the urgent need for restoration programs to explicitly consider biodiversity as a central objective^4^, and incorporate funding mechanisms based on robustly measured biodiversity outcomes^30^.

Funding based on carbon and biodiversity outcomes can take many forms in practice, including both the voluntary and compliance carbon markets (e.g., EU Emissions Trading System), and the nascent biodiversity credit markets^11,31^. The historical financial volatility of carbon markets, even within compliance markets, highlights the crucial role of public policy intervention. Mechanisms such as government-backed price floors and sovereign guarantees could establish a minimum guaranteed credit value, ensuring stable incentives for multi-objective restoration even during market fluctuations. At the same time, regulatory measures should follow clearly delimited rules and be independent from political interference to reduce risks of regulatory instability for investors^32^. Ensuring both price and regulatory stability helps nature credits to become credible instruments to provide carbon and biodiversity benefits.

The framework used in this study does not depend on any particular restoration funding mechanism. In practice, it is likely that a combination of public and private financial tools, including but not limited to nature credits, will be necessary to generate effective results^33,34^.

In our simulations, we define biodiversity credits based on the number of species with decreased extinction risks. We prioritize this species-based metric over area-based ones to better capture the “biodiversity uplift” central to emerging credit frameworks: “a measured and evidence-based unit of positive biodiversity outcome”^35^. Although our current analysis focuses on extinction risk, this framework is inherently flexible; it could be adapted to evaluate other metrics such as population recovery, geographic range expansion, or improved ecosystem services^36,37^. While a simulated setting allows us to quantify restoration at any level of demographic and spatial detail^38^, real-world implementation will require metrics that can be monitored accurately and cost-effectively, such as the periodic re-assessment of extinction risks for indicator species^39,40^.

A key distinction between carbon and biodiversity is that sequestered carbon is largely fungible –a ton of CO_2_ contributes similarly to climate mitigation regardless of where it is stored (barring issues of permanence^41^). In contrast, biodiversity outcomes are inherently context-dependent^36^; the value of recovering a specific species is tied to its rarity, its role in the ecosystem, and its geographic location. Estimating trade-offs among restoration outcomes and costs can help us define actionable and context-dependent valuation for *in situ* biodiversity credits, with their value likely to differ across geographic contexts, spatial extent, and the selection of reference species.

### Proof-of-concept application with real-world data

We demonstrated how our framework can be applied to real-world scenarios by analyzing a large empirical dataset from one of the most highly threatened and diverse ecosystems in the world, considered a Global Biodiversity Hotspot – the Atlantic Forest. Our analysis shows that our reinforcement learning approach can utilize diverse and multi-dimensional empirical data to identify priority areas for restoration, while balancing potentially conflicting targets and constraints for carbon sequestration, biodiversity outcomes, and implementation costs. While we focused here on the Atlantic Forest, the biodiversity simulation framework and optimization algorithms implemented in CAPTAIN can be adapted to different ecosystems and taxa and across spatial scales, through the parameterization of system-specific disturbance, recovery, and restoration actions. For instance, the framework has been applied at country or state levels in temperate and boreal forests^27,29^ and across ecosystems at the continental level^28^.

Within a 30% restoration scenario in the Atlantic Forest, we identified substantial discrepancies between carbon and biodiversity priorities, pointing to trade-offs that need to be accounted for to achieve positive change for climate and nature. Policies focused primarily on carbon sequestration tended to prioritize long and cohesive stretches of coastal areas (Fig. 4a). This encompassed most of the Brazilian coast within the tropical belt, probably due to the fast growth rates and larger carbon sequestering potential in those areas. In contrast, a biodiversity-only prioritization (Fig. 4a) identified a more scattered set of areas, probably due to the heterogeneous distribution of threatened species across the Atlantic Forest ecosystem.

Implementing spatial optimization tools in practice requires reliable ecological and socioeconomic data at a granularity consistent with the scale of restoration and conservation policies. While our analyses utilize empirical data, they rely on several simplified assumptions and approximations that must be addressed to transition from a proof-of-concept to real-world implementation. In particular, our use of land use as a proxy for anthropogenic disturbance and its modeled effect on plant diversity and abundance could be refined by characterizing disturbance patterns (severity, frequency, and projected trends) within each land use type using high-resolution data, and modeling their effects on plant diversity and abundance based on principles of plant community dynamics. Similarly, restoration costs –here approximated based on accessibility and land use– could be calibrated by empirical estimates of implementation and opportunity costs^42–45^.

Another challenge in real-world optimization of restoration strategies is quantifying the degree of uncertainty associated with the underlying data. While robust quantification of uncertainties exists or can be generated for some data (e.g., for climate projections and species distribution models^46,47^), equivalent estimates are not directly available for other data, such as human footprint and costs. Through our sensitivity tests we demonstrated that such uncertainties can be modeled through simulations, and that, as expected, they do influence the inferred restoration priorities (Fig. S9). As the amount and quality of spatial data continue to improve through data compilation, remote sensing, and emerging Earth foundation models^48^, it is likely that the availability of robust estimates of anthropogenic disturbance, costs, and other socio-economic variables will improve over time and allow increasingly detailed restoration planning. Potentially, the quantified uncertainties in restoration priorities can also be used to guide the acquisition of additional data targeting areas where new data are most likely to affect the prioritization outcomes.

The simulated restoration process through natural recolonization based on dispersal and growth could also be improved based on the findings of ongoing restoration experiments, to inform how different species can be re-introduced in restored habitats to accelerate the recovery of diversity and ecosystem functions^49–52^.

Crucially, the objectives, needs, and knowledge of Indigenous and local communities, which are essential for fair and effective conservation of nature^53^, should be integrated into the decision-making process. This could be achieved by restricting the range of candidate sites available for restoration to only comprise those that may indeed be available for that purpose, by imposing particular constraints on the optimization objectives deriving from the custodians of land, or by only including in the analyses species that are available and adequate for use in restoration. Work towards that goal is already underway in CAPTAIN^54^.

## Conclusion

The controlled simulations and empirical analyses performed here demonstrate the power and practical utility of a reinforcement learning framework to select priority areas for ecological restoration that substantially benefit climate and biodiversity. Focusing on a single restoration objective would result in missed opportunities^4^, whereas integrating both carbon capture and biodiversity conservation in a multi-objective prioritization allows us to predict the outcomes of different restoration policies.

Our model offers a way to explore how costs may be distributed between key financial mechanisms such as carbon credits and biodiversity credits^11^, which in turn can serve as a tool to estimate how a biodiversity crediting system could help achieve both climate and biodiversity targets. Future model developments could include additional funding sources, such as payments for ecosystem services (clean water, pollination) that increase over time as well-functioning ecosystems are recovered.

Ecosystem restoration is increasingly recognized as a crucial opportunity in the conservation of nature and stabilization of Earth’s climate^1,6^. The use of robust models and AI-based optimization, informed by the increasing availability of remote-sensed data and ground surveys, can help us deal with the complexity of biodiversity and its spatial and temporal dynamics, thus playing a critical role in guiding society’s efforts to tackle the twin crises of biodiversity loss and climate change.

## Methods

### Reinforcement learning optimization of restoration

We optimized spatial restoration planning by developing new functionalities within the CAPTAIN software^14^. We used environments and species generated through simulations to assess the performance and results of different restoration models under fully controlled settings. However, the framework can be used with empirical data, for instance by using species distribution models, climate data, and land use and other indexes of human footprint, and real costs of restoration. Similarly, empirical data, when available, can be used to inform the model about species dispersal ability, sensitivity to disturbances, and -in the case of plants- net sequestered carbon. In the following sections we first present how the updated CAPTAIN framework works for restoration with multi-objective reinforcement learning and then provide the details of how we set up the specific experiments presented in this study.

### Environment –

The environment is discretized into spatial units of arbitrary size that represent the cells of a grid and are used as units for the description of environmental variables, species ranges, and restoration actions. Each cell features environmental variables distinguishing land from sea and providing elevation and climate data (such as temperature and precipitation – e.g. their mean, monthly and seasonal variations; for ocean environments, specific variables such as water depth, oxygen levels, and salinity could be used). Cells are also characterized by a value of anthropogenic disturbance, indicated with *δ* ∈ [0, 1], that alters the ability of species to live in the cell and their abundance (see below) and by a restoration cost (*ε*), quantifying the relative cost of restoring the cell to a low level of disturbance. This could include, for instance, the cost of buying the cell, displacing or removing the cause of disturbance, and potentially taking action to reintroduce native species. Note that in real conservation work, restoration costs may be complex depending on the legal and socio-economic setting. For example, buying land is not always sufficient to stop all logging, but we assume that we can model appropriate total costs here. All these features can be set to change through time, e.g., capturing the dynamics of predicted climate change, variation in disturbance patterns, and evolving costs per cell.

The environment is also characterized by the species that live (or can live) in it. Specifically, for each cell and each species in the system, there is a value describing the number of individuals that can occupy the cell in the absence of disturbance (i.e. the natural state). This is a cell- and species-specific carrying capacity *K_c,s_* which depends on both the habitat suitability of the cell for that species (*H_c,s_*) and a species-specific maximum abundance per cell (*K_s_*):

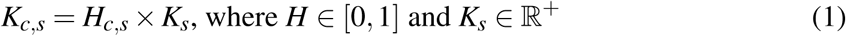

The environment also includes the actual number of individuals of a species at any given time *t*, *N_c,s,t_*, which results from the interaction between the species and the environment, including the effects of disturbance, growth rates, and dispersal.

For each species, we define a specific value of sensitivity *S_s_* ∈ [0, 1] that determines individual mortality fractions in response to a disturbance in a cell, *δ_c,t_*. Species are also assigned a growth rate *G_s_>* 1, which describes the demographic growth in one time step, and a dispersal rate *D_s_* ≥ 0, which determines the number of individuals that disperse from one cell to another. In the current implementation, we assume individuals to be static (e.g., plants). Therefore, dispersal does not move individuals from one cell to another. Instead, dispersal only adds individuals to the receiving cell (replicating for instance the dispersal of seeds). The number of individuals dispersing from cell *i* to *j* follows a truncated exponential distribution:

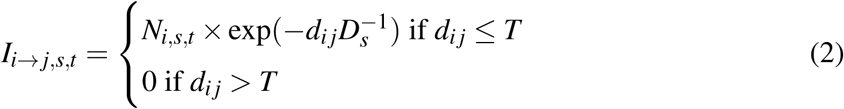

where *d_i j_*is the distance between the two cells and *T* is the maximum distance that can be reached in a single time step. Species distributions and abundances change over time based on growth, dispersal, and mortality. Specifically, at each time step, the distribution and abundance of species are updated according to the following equation:

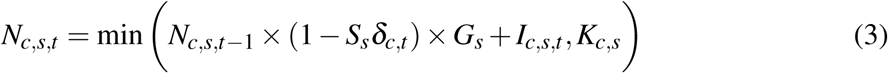

where *N_c,s,t_*_−1_ is the number of individuals of species *s* in cell *c* at the previous time step, *S_s_δ_c,t_* is the product between the disturbance in the cell at time t and the species sensitivity, with (1 − *S_s_δ_c,t_*) representing the mortality driven by disturbance. The term *G_s_* represents the contribution of demographic growth, while

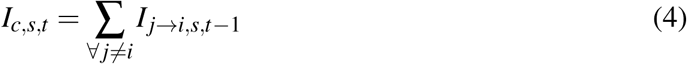

is the sum of the individuals dispersing from all other cells and represents the contribution of dispersal to the cell’s population.

Each species in the environment is assigned a value representing its expected carbon value *C_s_* ∈ R^+^. This is a single value and can be interpreted as the net amount of carbon sequestered by one individual of species *s*. As our simulated system in its current implementation does not track the age of each individual, we define the total carbon in a cell as the sum of carbon storage from all individuals and all species in the cell:

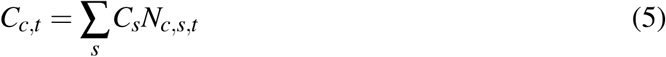

This quantity could in principle be extended to include carbon in the soil. We note that, as the age of individuals is not modeled here, the simulated accumulation of carbon in the current implementation might occur more rapidly than in empirical systems.

Finally, we also include a 5-tiered classification of species extinction risk, inspired by the International Union for Conservation of Nature (IUCN)^15^. When working on empirical data, this can be effectively replaced by actual IUCN Red List assessments or model-based predictions^40,55^. Hereafter, we will use the Red List-inspired labels to describe increasing extinction risks: Least Concern (LC), Near Threatened (NT), Vulnerable (VU), Endangered (EN), and Critically Endangered or (locally) extinct (CR). As part of the evaluation of the implemented restoration action, the initial extinction risk of a species can be updated based on the simulated dynamics of the species demography or geographic range, for instance, increasing or decreasing the extinction risk depending on whether the census population size of a species is decreasing or increasing, respectively. We indicate the extinction risk of each species *s* at time *t* as:

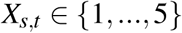

### Agent –

The agent in the system observes the environment at each time step extracting features for each grid cell about its disturbance level, its cost for restoration, the species that occur there, those that could occur there naturally in the absence of disturbance, their current extinction risk and carbon contribution (a detailed list of the features used in our experiments is provided in the next section). Following the modeling framework developed in the initial implementation of CAPTAIN^14^, the features of each cell feed into a fully connected neural network with tanh activation function^56^ in the hidden layers and a single output node. The parameters of the neural networks are shared across all cells and their outputs are pooled into a single vector of values providing a restoration rank of each cell. The agent will then select deterministically a predefined number of cells based on their ranking, within which restoration occurs. Here we assume restoration to result in the instantaneous removal of disturbance from the cells, although different dynamics such as a gradual decrease or partial reduction in disturbance can be implemented. The restoration action is associated with an economic cost computed as the cumulative cost of the selected cells. Once the disturbance is removed from a cell, its populations will continue to evolve according to Equation 3 with reduced (or removed) disturbance.

### Rewards and optimization –

The actions of the agent result in changes in the environment that are quantified through a *total reward*, defined based on the objective of the restoration policy. The parameters of the neural networks determining the restoration action are chosen to maximize the cumulative net reward at the end of a simulation of a fixed number of steps. The net reward or payoff is here defined as the weighted sum of three components quantifying net carbon gain (carbon reward), net biodiversity outcomes (biodiversity reward), and costs (budget reward). These quantities allow us to optimize policies with one or multiple objectives, generating socially optimal solutions when multiple objectives are used. At each time step, the carbon reward of a restoration action is computed as the ratio between the net change in total carbon stored in the system compared with its initial state (at time *t* = 0) and the amount of carbon lost between the natural state (in the absence of any human disturbance) and the initial state:

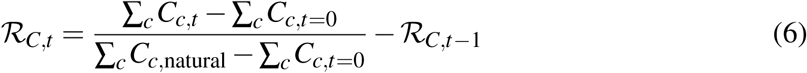

We note that with simulated datasets, like the ones used in this study, the natural carbon amount *C_c,_*_natural_, is known *a priori*. When working with empirical data, this quantity might have to be approximated based on measurements, such as through survey data (for trees, this can be derived indirectly from measurements of height and stem diameter at breast height (dbh), or more directly from biovolumes estimated from lidar scans).

The total reward of a restoration action also includes a measurement of the resulting biodiversity outcome, which can be defined in many different ways, as shown by a recent compendium^57^ currently listing over 570 biodiversity metrics. In the current implementation, we exemplify the calculation of biodiversity gains in a similar way as recently calculated in^38^, where the number of extinct or threatened species was used as a negative reward (see also^58^). The focus on extinction risk is also aligned with other recently proposed metrics for conservation priorities and threat abatement^39,59,60^. Here we use an average of the restored species population weighted by their current conservation status. Specifically, for any given extinction risk status *X* = *i*, we identify the subset of species *S_i_* such that:

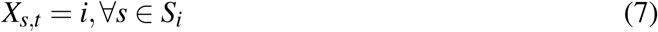

and compute the respective reward as

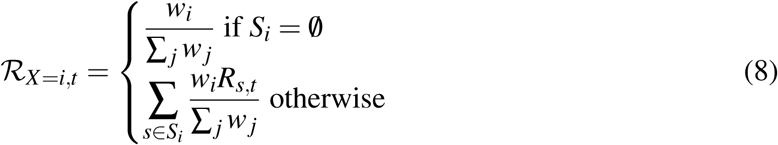

where *w_i_*is a weight associated with extinction risk status *i*, and *R_s,t_* is the ratio between the number of individuals of species *s* at time *t* found within restored cells (indicated with *M*) and the total population of the species in the environment:

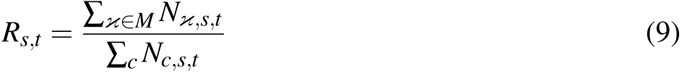

Thus, *R_s,t_*= 0 when the species is not found in any restored areas and *R_s,t_* = 1 when all its population falls inside restored areas. The weights are set to favor scenarios where the highest average fraction of population is restored in species classified in a high-risk category:

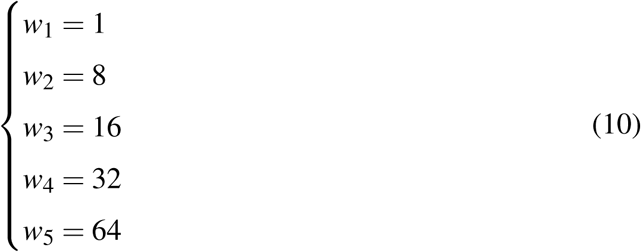

so that, e.g., protecting 30% of the individuals in a Least Concern species (*X* = 1) contributes 0.0025 to the reward, while the same fraction in a Critically Endangered species (*X* = 5) contributes 0.1587. The overall biodiversity reward is set to the sum of *R_X,t_* over all extinction risk categories. The use of increasing weights associated with extinction risk is consistent with several metrics developed to quantify conservation priorities^59,61,62^.

Finally, we include the restoration costs *ε_c_* as part of the reward. Based on an arbitrarily large reference budget *B_t_*_=0_ we compute the budget reward as

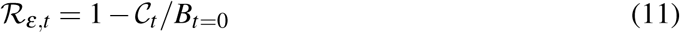

where C*_t_* is the total restoration cost summed over all restored cells (defined in the next section).

The rewards can be used alone or in combination to optimize policies that pursue one or more targets. Thus, we defined the total reward as:

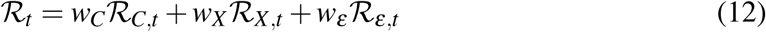

where the predefined weights (*w*) quantify the proportional importance assigned to carbon, biodiversity, and budget. Here we set the weight associated with the budget (*w_ε_*) to a constant and set the others to express their relative importance. Under this formulation, a policy targeting a single objective (e.g., focusing exclusively on budget) can be optimized by setting the respective weight to 1 and the others to 0 (e.g., *w_C_, w_X_* = 0, and *w_ε_* = 1). The weights can also be used to estimate different solutions where the total reward includes two or more potentially competing optimality criteria.

The CAPTAIN optimization algorithm trains a neural network policy to maximize the expected reward from protective actions by implementing a hybrid approach that combines parallelized evolution strategies (a genetic strategy algorithm) with concepts from classical policy gradient methods, i.e. an advantage function^14,63,64^. The algorithm operates through a repeated two-phase process (epochs): first, the policy parameters are randomly perturbed and evaluated via parallel environment simulation episodes; second, the results from these parallel runs are aggregated, and the parameters are updated using a stochastic gradient estimate. The parameter update is weighted by the advantage function, which measures the improvement of the current average reward relative to the running reward (weighted average of rewards across different epochs)^14^. This approach allows the policy to be optimized without the computational burden of calculating explicit gradients while enabling parallelization of the optimization task.

### Simulations

We use simulations of biodiversity to evaluate the ability of restoration policies optimized through reinforcement learning to achieve different targets, focusing on carbon and biodiversity outcomes. Although the same policies could be applied to empirical data, simulations allow us to benchmark and compare different restoration actions across many simulated scenarios in which we can accurately measure all outcomes.

#### Parameters of the simulated environment –

We simulated environments based on a raster of 128×128 cells, of 25 km resolution at the equator, inspired from physical properties of an arbitrary region spanning a portion of Central and South America (Fig. S2a). The properties include its coastline, elevation and cell-specific climate average values obtained from CHELSA^65^, here including the mean annual temperature, mean annual precipitation and the minimum temperature of the coldest month. We used the gdgtm Python library to generate the initial rasters (see Supplementary Data and Codes). The selection of variables is not [umath2]intended to provide an exhaustive account of the climate variables that determine a species’ niche but was used to generate artificial species distributions with realistically constrained geographic ranges. After *z*-transforming these topographic and climate variables we indicate them as *V* = {*V*_1_*, …,V_v_*}, where *V_i,x,y_* represents the value of the *i*^th^ variable corresponding to the longitude and latitude coordinates *x, y*.

We simulated species geographic range by first drawing a random pair of coordinates (*m, n*), based on which we extracted the corresponding elevation and climate data. We use a normal distribution to describe the environmental and climatic niche of the species MVN(*V_m,n_,* Σ), where the diagonal of the covariance matrix, a vector *σ* = {*σ*_1_*, …, σ_V_* }, is randomly drawn for each species from a uniform distribution U(0.05, 1) and off-diagonal values (covariances) are set to 0. The *σ* vector will therefore generate species with a range of narrow to wide climatic tolerances. We also introduce a distance factor, to ensure that species ranges are also spatially constrained e.g. through limited dispersal ability and not only determined by habitat suitability. The effect of distance was simulated through an exponential distribution with rate parameter *d* randomly drawn for each species from a uniform distribution U(0.05, 1). Thus, for any set of coordinates (*x, y*) the relative population density of a species is MVN(*V_x,y_,V_m,n_,* Σ) exp(−*d*Δ(*x, y, m, n*)), where Δ(·) is the Cartesian distance between (*m, n*) and (*x, y*). We use this formula to define the habitat suitability *H_c,s_*of a species *s* in each cell *c*, as used in Equation 1.

We used the empirical mean annual precipitation layer (rescaled between 0 and 1) to split cells into three categories with high (> 0.4), intermediate (0.25 – 0.4) and low precipitation (< 0.25). We use these categories to simulate different vegetation types, e.g. representing one dominated by trees, a mixed open habitat, and a fully open habitat (Fig. S2a). These vegetation types are then used to characterize properties of the species that inhabit them. Specifically, we simulated three categories of species to represent different growth forms, e.g. herbaceous plants, shrubs, trees. We assigned each species to one of the categories randomly with probability set equal to the proportion of the overlap of their simulated range with low, intermediate and high precipitation, respectively.

We drew the species demographic growth rates from a Weibull distribution:

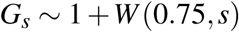

where *s* = 10, 2, and 0.1 for the first, second, and third growth forms, respectively. This results in faster average demographic growth for simulated ’herbaceous’ species. Similarly, we drew the species-specific carbon value from a gamma distribution:

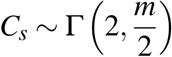

where *m* = 0.5, 1, or 2 is the mean of the distribution applied to the first, second, and third growth forms, respectively. We draw the species-specific carrying capacity *K_s_* (i.e. the maximum number of individuals that can live in a cell; Equation 1) from a beta distribution that generates many species with low abundance and few species with high abundance:

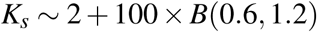

This leads to a mean carrying capacity of a species per cell of ≈ 35 ranging from ≈ 2 to ≈102. We assigned these values by setting the largest ones to species of the first growth form, the intermediate values to those of the second, and the smallest values to species of the third. Thus, species with fast growth and low carbon value tended to have higher carrying capacity than slow-growing species with high carbon value (reflecting the realistic differences in wood densities and carbon storage among woody species). Examples of the simulated species’ natural abundances resulting from both habitat suitability and species-specific carrying capacity are shown in Fig. S2d.

#### Disturbance and restoration costs –

We simulated spatially heterogeneous disturbance through random multivariate normal distributions rescaled to range between 0 and 1, which we multiplied by exp(-rescaled elevation) to simulate a scenario in which anthropogenic disturbance decreases on average with elevation (given the increased difficulties in accessing natural resources or performing agriculture; see example of simulated disturbance in Fig. S2c). We assigned a sensitivity to disturbance for each species based on a symmetric U-shaped beta distribution:

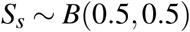

thus generating several species with either very low or very high sensitivity. Prior to applying any restoration policy, the environment was set to evolve for a number of steps until reaching an equilibrium under the simulated disturbance.

The restoration costs were linked to disturbance, so that on average the cost of restoration was higher for highly degraded cells (reflecting fewer opportunities for natural regeneration and higher distance to seed sources). We added some stochasticity in this pattern by computing costs as the product between disturbance and another rescaled random multivariate normal distribution (see example of simulated disturbance in Fig. S2c). We set the top 5% of cells with the highest cost as unrestorable, removing them from the set of cells that can be restored with an action from the agent. These cells are meant to represent areas where restoration is not possible e.g. because that would imply displacing inhabited areas or activities and infrastructure that cannot be moved.

#### Evolution of the species extinction risk –

We initialized the species extinction risk based on their range size, on their sensitivity, and on the effect of disturbance:

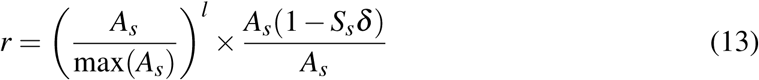

where *A_s_* is the natural range of the species (number of occupied cells) in the absence of disturbance. Thus, the value *r* decreases with small natural range sizes and with increasing sensitivity and/or disturbance^6^. The exponent *l* (here set to *l* = 0.3) has the effect of flattening the difference in initial extinction risk based on natural range size. We then mapped the value to an initial extinction risk *X_s,t_*_=0_ (Table 1). We note that this approach to simulate species extinction risk, while based on common assumptions about species range size and decline due to anthropogenic disturbance, is arbitrary and should be replaced with actual risk assessments (e.g. from the IUCN Red List) when running the optimization with real world data.

Throughout the simulations, the extinction risks of all species were re-evaluated at each step, and we used two criteria to re-assess the conservation status of a species. First, we calculated the number of restored cells within which a species *s* occurred at time *t*, indicated as *O_s,t_* ∩ *M_t_* relative to the species natural range size *A_s_*. Thus, increasing values of (*O_s,t_* ∩ *M_t_*)*/A_s_* indicate that a proportionally higher number of restored cells is occupied (potentially after re-colonization) by species *s*, leading to a decreased extinction risk (Table 1, Criterion 1). Second, we calculated the ratio between the occupied geographic range at time *t* and the occupied range at the beginning of the simulation *O_s,t_*_=0_, i.e. prior to enforcing restoration programs. This ratio is used to verify if the species is in decline, e.g., if the restoration is ineffective or if other factors, such as climate change, negatively affect a species’ range. We used the thresholds reported in Table 1 (Criterion 2) to determine if the species was in decline and to what extent that affected the species extinction risk. Finally, any species among the initial simulated pool with occupied range size of 0 was moved to the highest risk category and considered as possibly extinct in the study system. These criteria are inspired by some of the Red List guidelines^66^, specifically looking at species range sizes, and recovery or declining trends through time. While we used the occupied range as the main metric, the criteria can be defined based on alternative metrics, e.g., based on census population size and its temporal variation or fragmentation of the species range.

#### Observed features –

The agent was set to observe the entire environment at each step, extracting several features for each cell, which form the input of the neural network and determine the next action. We used the same 16 features for all policies, independently of the reward. The first features were the disturbance in the cell and the mean disturbance in the 5×5 cells around it (features 1-2). The latter feature provides a measure of the spatial distribution of disturbance, for instance, differentiating between a disturbed cell neighboring a pristine area and one placed within a broader degraded area. We then included the potential carbon net change in the cell calculated as the difference between the total carbon in the cell under natural conditions and the current amount of carbon, i.e. *C_c,_*_natural_ − *C_c,t_*. The same measure is also provided as a mean value in the 5×5 cells around the cell (features 3-4). The features additionally include the number of CR, EN, VU and NT species in the cell based on the current populations occupying the cells and the mean number of CR and EN species in the surrounding 5×5 cells (features 5-10). We also included the number of species in each of the five extinction risk categories, but counting only species that do not occur in any of the previously restored cells, if any (features 11-15). These features serve as a measure of complementarity among cells so that the model can choose to restore cells with a different species composition compared with areas already restored. Finally, we included the cell’s cost (feature 16). All features were rescaled between 0 and 1 before being fed to the neural network.

#### Training –

We trained models with 250 species running simulations for 30 time steps (such as years). The agent was set to restore 100 cells at each step, reaching 3,000 restored cells at the end of the simulation, i.e., corresponding to about 30% of the land cells in our grid to mimic the 30% restoration target of the Kunming-Montreal Global Biodiversity Framework. We used a parallel genetic strategies algorithm with 8 environments evolving in parallel and ran 5,000 epochs to train the models. We monitored the reward across epochs to assess model convergence (Fig. S4). We used small neural networks for each cell, connecting the 16 features to a first hidden layer of 3 nodes, followed by a second hidden layer of 2 nodes. The parameters of the network were shared among all cells. We also tested a larger neural network model, which however did not lead to significant performance improvements in our experiments (see *Supplementary Information; Fig. S5*). We trained different models exploring a gradient of optimality^26^ going from favoring the carbon outcome to favoring the biodiversity outcome. We achieved this by assigning different weights to the reward (see Equation 12). Specifically, we trained five models, keeping the weight of the budget reward constant at 0.5 and assigning different weights to carbon and biodiversity such that the sum of the weights was kept constant and equal to 1 (Table 2).

**Table 2:**
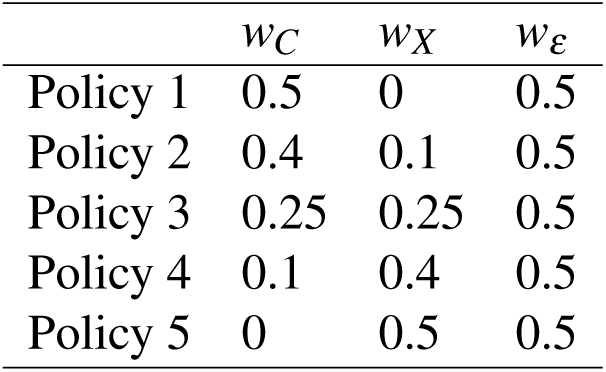
Weights used in the different trained models and assigned to the carbon reward (*w_C_*), biodiversity reward (*w_X_*), and budget reward (*w_ε_*). We considered a range of models keeping the weight on the budget reward constant, while progressively moving from a carbon-focused optimization (Policy 1) to a biodiversity-focused one (Policy 5). The intermediate policies allowed us to explore the trade-offs between these potentially competing objectives.

#### Experiments on the reference region –

After training the models, we used them on 100 simulations each including 500 species and initialized following the same approach utilized to generate the training data. These simulations were run on the same reference region used for the optimization of the policies. Although these simulations were based on the same simulation settings, they differed from each other, being based on random draws for all parameters. Thus, the distribution of species, disturbance, and costs and the traits associated with each species (carbon, sensitivity, growth rates) varied randomly across simulations. This allowed us to evaluate the results of different models across a range of different parameter values. The simulated environments in their initial state, i.e. prior to restoration, included on average 30 CR species (standard deviation, sd: 18), 87 EN (sd: 39), 156 VU (sd: 31), 139 NT (sd: 30), and 87 LC (sd: 56) species. As during training, the simulated datasets evolved for 30 time-steps with the agent interacting with the environment based on each of the five trained policies and restoring 3,000 cells. We evaluated the outcomes of the different policies by calculating the amount of carbon gained through restoration, the changes achieved in terms of number of species falling in each of the five extinction risk categories, the number of species occurring within restored areas at the end of the simulation, and the total cost of restoration. This allowed us to quantify the trade-off between carbon and biodiversity outcomes along a gradient of optimal solutions.

### Carbon and biodiversity credits

We implemented a simplified concept of carbon credits by assigning a monetary value to the amount of net carbon gain (or potentially loss) incurred during the simulations. As carbon (like restoration costs and budget) is set in arbitrary units in our simulations, we defined the value of a carbon credit unit (*χ*) as the value of the net carbon gain obtained through Policy 1, namely the policy placing the highest weight on carbon, leading to the full compensation of the restoration costs:

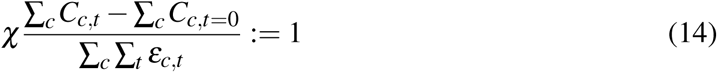

where *χ* is the value of one carbon credit unit based on the simulated systems used for carbon and budget, the fraction’s numerator is the net carbon gain at time *t* compared to the initial state (*t* = 0) and the denominator is the total cost of restoration. Thus, under this definition, the carbon credits accumulated through the implementation of Policy 1 result, by construction, in full compensation of the cumulative restoration costs, with no economic loss at the end of the simulation. We note that this definition is arbitrary and could be replaced by other definitions, including empirical valuations of carbon in the cap-and-trade system of the European Union, or others.

The same carbon credit value *χ* was then applied to the other four policies where carbon was given lower weights (Policies 2–5) in the training phase with the inclusion of a positive weight assigned to the biodiversity reward. This allowed us to evaluate to what extent the carbon credits accumulated through restoration policies that optimized for both carbon and biodiversity (Policies 2–4) or biodiversity only (Policy 5) were able to compensate for the restoration costs.

Next, we selected a simple metric as a reference to define a biodiversity credit and developed an approach to value the credit relative to the value of a carbon credit. Specifically, we quantified the biodiversity credits as the number of species that moved from a threatened class (CR, EN, VU) at the beginning of the simulation to a non-threatened class (NT, LC) after restoration. This differs from current practice in some of the emerging biodiversity credit markets, such as in Colombia where one biodiversity credit unit equals one hectare regardless of how many species benefited from the restoration and in which ways. However, the same approach utilized here can be used for alternative definitions of biodiversity credits.

We then evaluated the net financial loss or gain of each policy after stacking the value of carbon and biodiversity credits under different valuations of biodiversity credits. We indicate with *β* the value of one biodiversity credit obtained from moving a species from a threatened to a non-threatened status after restoration. Given that we are using arbitrary units for costs and carbon credits, we defined the value of a biodiversity credit relative to that of a carbon credit. This allowed us to assess which policy would become financially viable or preferable, in our simulated context, if e.g. the value of one biodiversity credit equals that of a carbon credit (i.e. *β* = *χ*). We tested valuations of *β* = 0*, χ,* 2*χ,* and 4*χ* and calculated the net financial loss or gain of a policy (Φ) as a percentage of the restoration cost:

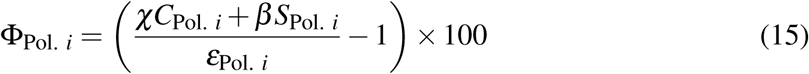

where *C*_Pol.*i*_ is the net carbon gain of the *i*^th^ policy, *S*_Pol.*i*_ is the policy’s net biodiversity outcome (as defined above), and *ε*_Pol.*i*_ is the cost of restoration.

Finally, we calculated, for any given policy *i*, the value that should be attributed to a biodiversity credit so that the policy results in the same net financial outcome as a restoration program focused on carbon (i.e. our Policy 1):

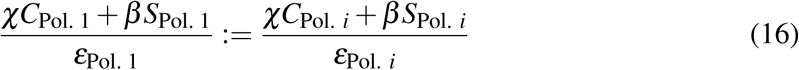

The value of one biodiversity credit is therefore, after solving Equation 16 for *β* :

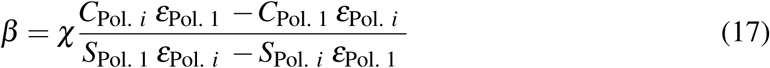

Using this formula, we derive the value of a biodiversity credit (relative to the value of a carbon credit) required to make the financial outcome (defined in Eq. 15) of any optimal restoration policy equal to the profit of a carbon-oriented policy (our Policy 1), that is, such that Φ_Pol._ *_i_* = Φ_Pol. 1_. We note that under this definition, the value of biodiversity credits is context-specific, as it depends on the overall area considered and its restorable carbon and biodiversity. Within the area, the restoration of cells that contribute the most to reducing the extinction risk of species (e.g., cells that protect rare and threatened species) will lead to the highest biodiversity credits^67^.

### Empirical analysis

We analyzed an extensive dataset of tree biodiversity in the Atlantic Forest^19^. The dataset included occurrence data for thousands of woody plant species in the Atlantic Forest, with per-species information on their maximum height, wood density, and conservation status based on the IUCN Red List classification of extinction risks^15,19^.

We compiled spatial data gridded into 16,376 cells at a 10-km resolution within the Atlantic Forest region (around 1.2 × 10^6^ km^2^ in area) as defined by the WWF Global 200 project^68^. We generated species distribution models (SDM) based on cell-specific WorldClim bioclimatic variables^69^ and on species occurrence data^19^, and generated raster layers representing disturbance and restoration costs based on publicly available geospatial data products. All data processing and analyses were run in R 4.3.3, using the packages concaveman, ENMTools, flexsdm, future, fuzzyjoin, geodata, magrittr, sf, terra, tidyterra, and tidyverse^70–81^.

#### Species distribution models –

We obtained the 19 WorldClim historical bioclimatic variables at 5 arc-minutes resolution^69^, cropped to the Atlantic Forest region and reprojected to the 10- km resolution. We examined the correlation between the bioclimatic variables to exclude highly collinear variable pairs, retaining the following 7 variables of temperature (^◦^C) and precipitation (mm): 1) annual mean temperature, 2) mean diurnal temperature range (mean of monthly maximum temperature - minimum temperature), 3) annual temperature range, 4) annual precipitation, 5) seasonality of precipitation (coefficient of variation), 6) precipitation of the warmest quarter, and 7) precipitation of the coldest quarter.

Among the retained variables, only variable pairs 1-5 (*r* = 0.65) and 2-7 (*r* = 0.61) exhibited coefficients of correlation larger than 0.6 (Fig. S10): as these are both pairs of a temperature variable and a precipitation variable and they may contain important environmental information despite the correlation, we retained both variable pairs.

We obtained geolocated occurrence data for 4,950 species occurring in the Atlantic Forest^19^, and eliminated data points outside of the rectangular bounding box of the Atlantic Forest region to exclude geographically distant data points. The remaining data set contained in total 722,448 records and 4,948 species. Of these, 3,105 species had abundant data points (*n* ≥ 30) and 1,631 species had sparse data points (3 ≤ *n <* 30); 212 species with insufficient data points (*n <* 3) were excluded from the analysis. To mitigate potential spatial biases, we performed spatial thinning for species with abundant data points, using the ENMTools::trimdupes.by.raster function to retain one data point at maximum per pixel. To prevent excessive loss of species information, we only accepted the thinned results if they also contained ≥ 30 data points; otherwise, we retained the original unthinned data. In total, 1,843 species were thinned, and for the thinned species 80% of data points were retained per species on average.

We defined the spatial extent of the training area (from which background points are sampled) for each species as an alpha shape (concavity = 3) surrounding all occurrence data points of the species, buffered by twice the median inter-point distance among all data points and cropped to the Atlantic Forest region. We sampled 10,000 background points within the training extent of each species^82^, assembled the occurrence data points and the background points, and extracted the environmental values at all points to construct the model data. We partitioned model data into training and testing sets for cross-validation, using complete random *k*-fold partition implemented in the flexsdm::part_random function: we used 4-fold partition for the 3,105 species with abundant data points, and 2-fold partition for the 1631 species with sparse data points.

We then fitted an SDM for each species: for the 3,781 species with ≥ 15 data points, we fitted a generalized additive model (GAM) using the flexsdm::fit_gam function, and for the 955 species with *<* 15 data points, we adopted an ensemble of small models (ESM) approach using the flexsdm::esm_gam function, which consists of fitting bivariate GAM models with all pair-wise combinations of predictors, and calculating the ensemble average of model outputs weighted by their performance metrics (Somers’ *D*: *D* = 2 × (AUC − 0.5))^83^. For the model output of each species, we calculated the following five performance metrics: sensitivity, specificity, omission rate, true skill statistic (TSS), and area under curve (AUC). We excluded species with low performance (sensitivity *<* 0.7, specificity *<* 0.7, TSS *<* 0.5, AUC *<* 0.8), retaining results for 3,885 species (Figure S11). Finally, for the retained species, we generated geographical predictions of continuous suitability score maps over the Atlantic Forest region using the flexsdm::sdm_predict function.

#### Disturbance and restoration costs –

We defined the disturbance level within each 10- km pixel of the Atlantic Forest region based on its land cover. We obtained the MapBiomas Collection 10 - Land Cover and Use dataset^84^, which classifies each 30-m pixel within Brazil into one of 38 land cover classes. In addition, we also obtained the MapBiomas Collection 9 - Age of Secondary Vegetation maps in 2023^85^, and created a mask of secondary forests representing pixels where the age of secondary vegetation *>* 0. We cropped the datasets to the Atlantic Forest region, reprojected to the 10-km resolution, and calculated the number of 30-m pixels belonging to each land cover class within each 10-km pixel, additionally classifying pixels within the secondary forest mask into a novel class of “secondary forest” (ID: 76). We reclassified the MapBiomas classes into the following 9 broad classes:

- Forest (IDs: 1, 3, 5, 6, 49),
- Grassland (4, 12, 29, 50),
- Secondary forest (76),
- Plantation (9),
- Cultivation (14, 15, 18, 19, 20, 21, 31, 35, 36, 39, 40, 41, 46, 47, 48, 62),
- Coastal area (23),
- Water bodies (11, 32, 26, 33),
- Urban area (24, 25, 30, 75), and
- Others.

We calculated the proportion of each broad class within each 10-km pixel. We then calculated the average disturbance score of each 10-km pixel by assigning a disturbance score to each broad class, namely, forest and grassland: 0, secondary forest: 0.3, plantation: 0.5, cultivation: 0.7, coastal area: 0.5, water bodies: 1, urban area: 1, others: 0.5. We then calculated the weighted average of all disturbance scores weighted by the proportion of the broad classes.

We estimated restoration costs by integrating three spatial proxies: (1) accessibility (travel time to urban centers) and (2) elevation, which serve as proxies for logistical and implementation challenges, and (3) agricultural production value, representing the opportunity cost of land conversion. While this approach provides a robust approximation for our case study, we acknowledge that accurate restoration costs would require more granular data, including the costs of active reforestation, fencing, invasive species removal, and long-term site management^6,42,86^.

- For accessibility, we retrieved data of travel time to cities with more than 5,000 inhabitants (geodata::travel_time)^87^.
- For elevation, we retrieved global elevation data (geodata::elevation_global) that are based on the Shuttle Radar Topography Mission (SRTM) 90-m resolution data^88^ and supplemented by the GTOPO30 dataset for high latitudes^89^.
- For the value of agricultural production, we retrieved SPAM data of aggregated crop production value (USD) per area (geodata::crop_spam).

SPAM pixels with empty values were filled in by interpolation from edges using the GDAL script gdal_fillnodata, with the md (maximum number of pixels to be searched for values to interpolate) set to 10. All three data layers were cropped and reprojected using the GDAL command gdalwarp, resampled to the 10-km resolution with bilinear interpolation using the terra::resample function, and then scaled with min-max scaling to the range of [0, 1]. We then calculated restoration costs as the arithmetic average of the three scaled variables.

#### Species traits –

We included species-specific features in our analysis, allowing us to incorporate extinction risks, proxies for carbon sequestration potential, and life-history traits. From the original data^19^ we obtained growth habit (categorized as shrub, small tree, and large tree), which we used to approximate relative values of growth rates. We used the same Weibull distribution used in the simulated data to draw species-specific growth rates, with scale parameters set to 10, 2, and 0.1 for shrubs, small trees, and large trees, respectively. This led to faster average demographic growth rates for shrubs compared to large trees. We acknowledge that these are approximations and it would be preferable to replace this parameterization with empirical estimates of growth rates, where possible, for instance leveraging increasingly comprehensive databases such as TRY^90^.

We randomly drew the values describing species-specific sensitivity to disturbance from the same U-shaped beta distribution used in our previous simulations. To simulate the fact that trees might be more sensitive to land use changes, we ranked the randomly drawn sensitivity values and assigned the highest values to large trees, followed by small trees, leaving the smallest values to shrubs. Again, these approximations should be replaced by empirical estimates of sensitivity to disturbance, where available. We expressed the carrying capacity of a species in a cell as a relative number between 0 and 100, scaled by the habitat suitability in the cell, as predicted by the species distribution models. The simulated abundance of a species in a cell was further affected by a mortality rate proportional to the product between disturbance and species sensitivity (Eq. 3).

We used the available Red List assessments as species extinction risk estimation^15^. Here we imputed missing extinction risk classification (i.e. unevaluated or data-deficient species) through random resampling proportionally to the empirical frequencies of each class among assessed species. While this is a crude approximation, more sophisticated approaches could be used to predict extinction risks using machine learning imputation methods^55,91^. We let the extinction risk of species evolve over time as a function of species recovery after restoration using the re-assessment criteria used in our simulations (Table 1).

We used per-species biomass as a proxy for a species’ ability to sequester carbon, based on wood density (WD) and maximum height (H)^19^. We estimated the maximum diameter at breast height (DBH) for each species from maximum height (H), using the generic model 3 from^92^: log(H) = 1.029 + 0.567 × log(DBH) from which we obtain DBH = exp((log(H) − 1.029)*/*0.567). Next we estimated maximum above-ground biomass (AGB, kg) per tree for each species from wood density (WD), DBH and H, using the pantropical model from^93^: AGB = exp(−2.977 +log(WD×DBH^2^ ×H)) ≈ 0.0509 ×WD×DBH^2^ ×H. As in the analysis of simulated data, we estimated the relative above-ground carbon in a cell based on the species-specific AGB estimated as shown above and the simulated abundance of the species in the cells. We then calibrated this relative quantification of carbon per cell, by rescaling it such that their average equaled 98.5 tonnes per hectare, based on previous empirical estimates for the Atlantic Forest^21^. We then used the difference between the initial carbon and the total carbon after restoration as a measure of the sequestration generated by the restoration policy. While these are approximate estimates of potential carbon heterogeneities in the region based on species composition, advances in above-ground carbon estimation for instance from remote sensing data^94,95^ will likely improve the quality of the restoration prioritization in the future. For example, high-resolution remote sensing data products^96^, in combination with ground data such as forest inventory plots^97^ and terrestrial laser scanning^98^ in areas where they are available, could be used to provide more accurate estimates of the species-specific carbon sequestration potential.

#### Model fine-tuning and predictions –

We fine-tuned the models by running another training procedure for 1100–1700 epochs starting from the parameter values from the models trained on simulated data (Policies 1, 3, and 5). The additional training was based on a batch of 6 parallel environments, each containing a random subset of 1,000 species to speed up the run time.

We then used the fine-tuned model on the full set of species to identify restoration priorities. As in the analyses of simulated data, we set the models to select 4,800 grid cells (i.e. approximately 30%) within 30 time steps. Thus, at each time step 160 cells were selected for restoration, and the disturbance in protected cells was set to 0, allowing existing species in the cell to reach their carrying capacity and species from neighboring cells to recolonize them based on habitat suitability and a dispersal process modeled as in our previous simulations.

As some of the parameters in our simulations were imputed (namely, the extinction risks for un-assessed species) or randomly drawn from prior distributions (i.e., the growth rates, and sensitivity to disturbance), we repeated the restoration predictions 30 times, resampling each time these parameters. We then summarized the results by ranking cells based on how frequently they were selected for restoration and took the top 4,800 cells as the final outcome. We additionally performed sensitivity analyses where we introduced random variation in the carbon and cost estimates to assess the impact of error in our estimates on the predicted restoration priorities and outcomes (*Supplementary Methods and Results* and Fig. S9).

## Data and code availability

All the code and data used to reproduce the results shown in this study will be available on Dryad. The CAPTAIN software is available here: https://www.captain-project.net

## Generative AI statement

Generative AI was not used in producing the contents of this article.

## Supplementary Information

### Supplementary Methods and Results

#### Generality of carbon and biodiversity trade-offs

We tested to what extent our policies trained on a single arbitrary region could be applied to other geographic and climatic contexts, by running restoration experiments on a set of 100 additional datasets of varying geographic location, extent, and resolution (Fig. S3). Specifically, we generated 100 additional rasters of 128×128 cells with resolution at the equator of 2.5, 10, or 25 km. Their placement globally was random but conditioned on having at least 70% of cells on land (Fig. S3). For each raster we extracted elevation and climate data and simulated 500 species following the procedure described above. We then ran the five optimized policies on these datasets to evaluate the ability of our trained models to generalize outside of the region on which they were optimized.

We found that, despite such wide geographic variation and no fine-tuning to regional conditions, the policies consistently resulted in general trends reflecting the objectives used to train them. Accordingly, the net carbon gain decreased from Policy 1 to 5, the number of species with improved conservation status was higher in Policies 2–5, and costs were roughly constant across all policies (Fig. S7a–c). However, the patterns were less consistent with regards to changes in species number across risk categories (Fig. S7d–i). These results suggest that while models trained on a region can be applied with reasonable performance to other regions, they would almost certainly benefit from a region-specific re-training or fine-tuning.

Consistent with our previous experiments, we found that the restoration areas selected under different policies only partly overlap (Fig. S1b), indicating that trade-offs between carbon and biodiversity priorities are common at different geographic locations and scales. These experiments also confirmed the crucial potential of biodiversity credits in contributing to the financial feasibility of restoration policies that target both carbon and extinction risk reduction (Fig. S8a–d; Table S2).

#### Comparison with greedy search optimization

We carried out analyses of the simulated datasets using simple metrics to select the restoration areas instead of CAPTAIN’s neural network to evaluate the performance of our trained models against simpler non-parametric approaches. We ran three models targeting carbon and cost, extinction risk and cost, or all three factors. The models used a subset of the 16 features monitored by the agent in the other experiments.

Non-parametric model (NPM) 1 focused on carbon and costs and used ratio between the potential carbon in a cell (feature 3 in our setup; see *Methods, Observed features*) and its restoration cost (feature 16) as the priority metric to select restored areas. NPM 2 targeted carbon, extinction risk, and costs. We summarized the importance of a cell for biodiversity as the weighted average of the number of CR, EN, VU, NT, and LC species (all rescaled between 0 and 1). To maximize the coverage of protected species we only counted LC species that are not already found within previously restored cells (feature 15 in our CAPTAIN policies). The weights were proportional to those used in our CAPTAIN reward function (Eq. 10). Restoration priorities were set proportional to the ratio between the sum of carbon and biodiversity metrics and costs. Finally, NPM 3 focused on extinction risk and costs and selected the cells with highest ratio between the biodiversity metric and costs.

Through the analysis of 100 simulated datasets we found that NPM1 achieved a performance similar to that of the equivalent neural network model (Policy 1) with comparable net carbon gains and slightly lower costs (Table S3). The biodiversity outcomes of NPM2-3 (i.e. models that included extinction risk reduction in their objectives) were however generally worse than those obtained from trained policies. For instance NPM3 (which did not include a carbon objective) included on average 32 fewer species in the selected areas compared to the equivalent trained model (Policy 5). It also led to the recovery of 133 previously threatened species as opposed to the more than 200 species achieved by trained models. The restoration costs of NPM2-3 were also slightly higher than those of equivalent trained policies (Table S3). This shows that the additional computational burden of training a neural network policy is counterbalanced by an improved outcome.

#### Testing a larger neural network

We compared the performance of the small neural network utilized in our CAPTAIN experiments against that of a larger model. To this aim we trained additional models with two hidden layers of 16 and 8 nodes, using the same training procedure and simulated datasets (Fig. S5). These models therefore included about 7 times more parameters than the small ones.

After running the trained models on our set of 100 simulations, we found that the larger models did not yield substantially improved outcomes compared to the small models (Table S3). Costs, carbon and biodiversity metrics suggest that the richer parameterization does not lead to a significant change in performance under our simulated scenarios. These results are likely a function of the relatively small number of features used in our analyses and larger models might be beneficial in scenarios with more aspects of the environment being observed by the agent. Although the construction of the model with parameter-sharing across grid cells and rescaled features makes the number of parameters independent of the size of the area and the number of species, further exploration is needed to assess whether larger models might lead to a better performance with datasets spanning a wider extent and more complex biodiversity patterns.

#### Sensitivity analyses for the Atlantic Forest restoration priorities

We ran additional analyses to explore the potential impact of inherent error in the input data, focusing on the variables which we deemed most likely to influence the results: the species-specific above ground biomass (AGB; used as a proxy for carbon sequestration) and on the spatial distribution of restoration costs. While we acknowledge that these experiments do not capture all sources of uncertainty and error in the data, they serve as a demonstration of how the robustness and sensitivity of estimated restoration priorities can be assessed.

#### Simulating error in carbon estimates –

First, we simulated error in AGB by randomly drawing for each species *i*,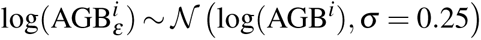. This resulted in a simulated error of ≈ 20% on the carbon associated with each species relative to our estimates. We performed 30 randomizations of the species AGB values and analyzed them under the three trained policies.

#### Simulating error in cost distribution –

We additionally simulated error in cost distribution using Perlin noise to generate random, but spatially structured variation (Fig. S13 for an example). The random noise was centered in 1, rescaled to range from 0.75 to 1.25, and applied as a multiplier to the cost layer (Fig. S12a). This resulted in a ± 25% random variation of the costs. We performed 30 randomizations of the cost layer and analyzed them under the three trained policies.

Our analyses show that, although the prioritization maps remain broadly consistent among experiments, the additional error has a non-negligible impact on the inferred restoration priorities (Fig. S9). We measured the effect by computing the average percentage overlap between the priorities inferred from each of the 30 replicates and those identified in the absence of additional simulated error (i.e. the area shown in Fig. 4).

The average overlap across runs with error in carbon estimates was 94% for Policy 1, 79% for Policy 3, and 66% for Policy 5. The overlap decreased in the presence of spatially structured error in cost estimates, to 87% for Policy 1, 69% for Policy 3, and 51% for Policy 5. These results show that, while most of the priority areas are consistent across sensitivity tests, errors in the data can lead to more uncertain predictions, with a greater impact on restoration priorities targeting extinction risk compared to carbon (Fig. S9f, i). Our analyses show that this approach can be used to assess the sensitivity of prioritization results to data uncertainties and identify top-priority areas that are robust to error in the data, while highlighting the areas that are most prone to shifting priority.

## Supplementary Figures

**Figure S1:**
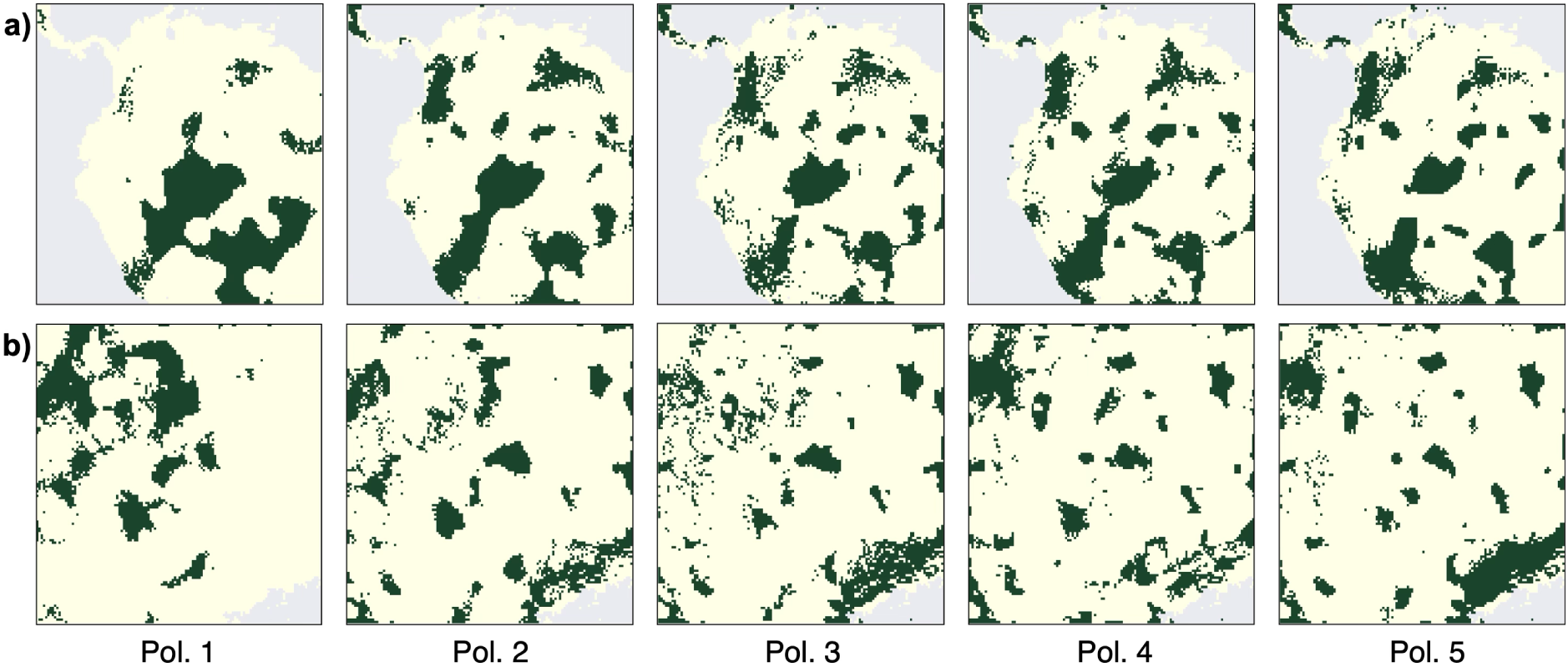
Changes in spatial restoration priorities depending on the relative weights of carbon (highest in Policy 1) and biodiversity (highest in Policy 5). a) Example of the restored areas in one of the 100 simulations performed on the reference region used to optimize the models (Fig. S2) selected under five policies (Table 2). Note that the placement of the restored areas depends on the specific conditions of the simulated dataset. Restored areas will therefore vary among simulations, reflecting the randomly initialized distribution of species, disturbance, and costs. b) Inferred restoration priorities obtained from simulations based on another, randomly selected region, showing similar discrepancies in spatial prioritization depending on the relative importance attributed to carbon vs biodiversity outcomes (Fig. S3).

**Figure S2:**
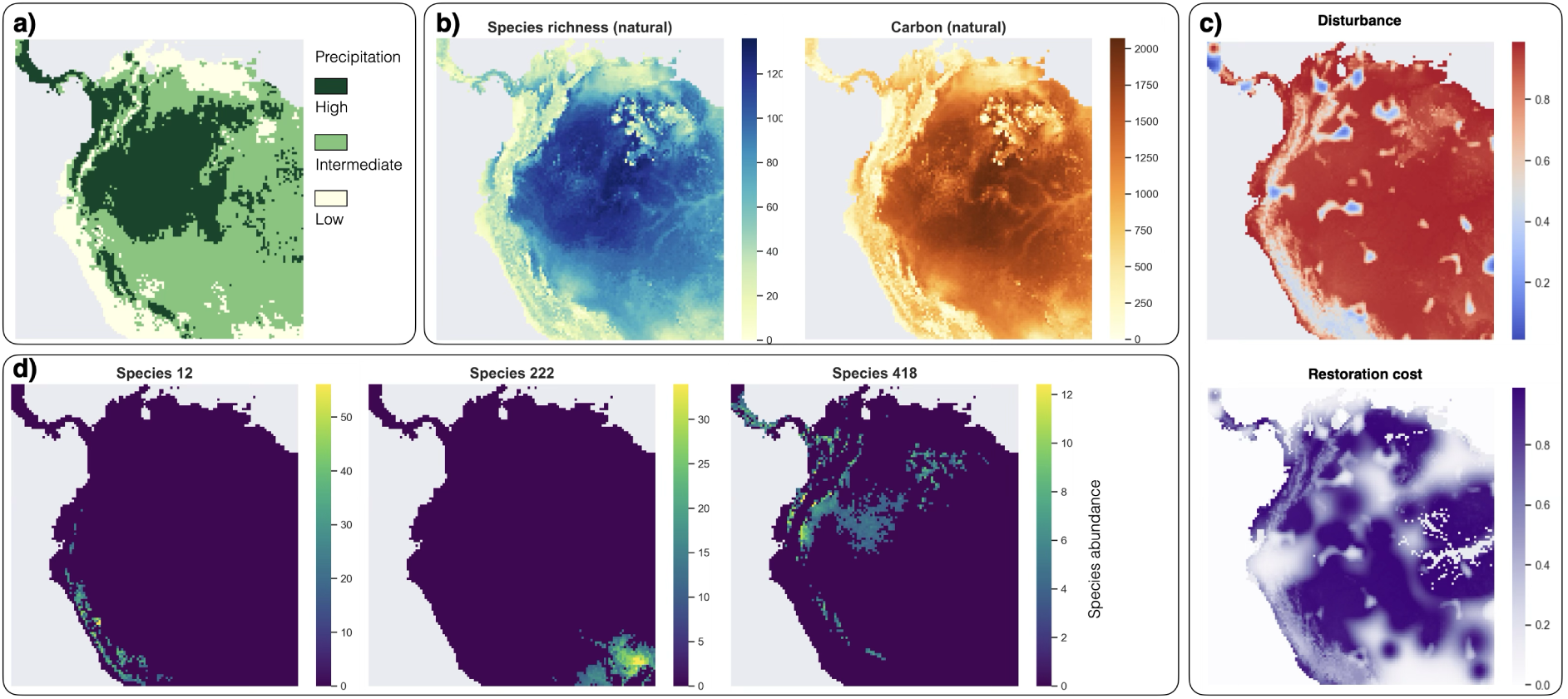
Map of the arbitrary area used for model training and subsequent simulations with examples of the empirical and simulated data used in our CAPTAIN runs. The physical properties of the area are based on the real topography and climate of the selected portion of Central and South America (a). The area was gridded into 128×128 cells and included empirical climate and elevation layers, while the ’natural’ state of biodiversity and carbon (b) and the disturbance patterns and costs (c) were simulated and changed randomly across simulations. Species richness and carbon maps are obtained after stacking 500 simulated species (see examples in d). Note that only the map in panel a) is based on real data, while all other panels show simulated data. Similar simulations were repeated on 100 additional random maps of different size and location (Fig. S3).

**Figure S3:**
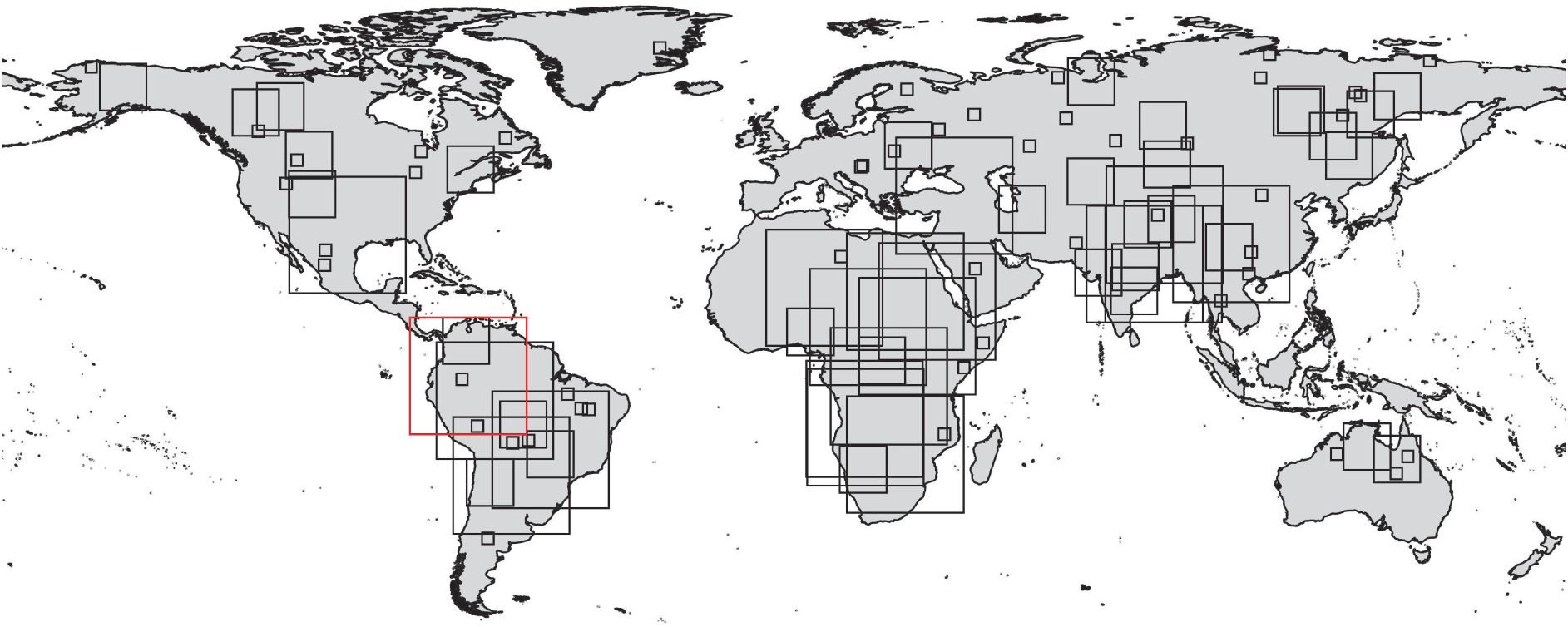
Maps showing the 100 randomly sampled rasters used as the basis for our simulations. The raster highlighted in red indicates the region used to train the models and to run the simulations based on a fixed geographic extent (see Fig. S2 and *Methods*).

**Figure S4:**
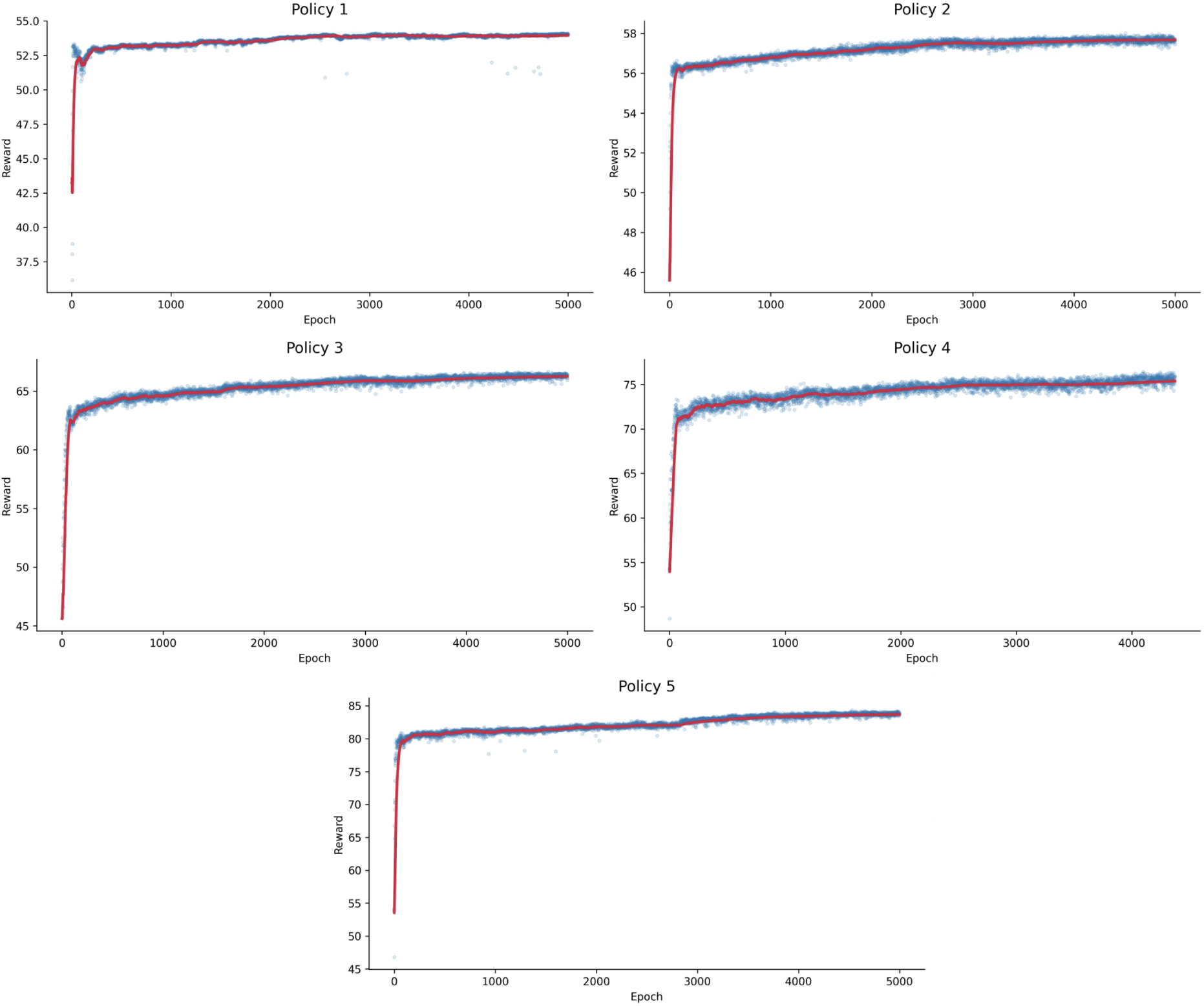
Training rewards across epochs based on a small policy network (see *Methods*).

**Figure S5:**
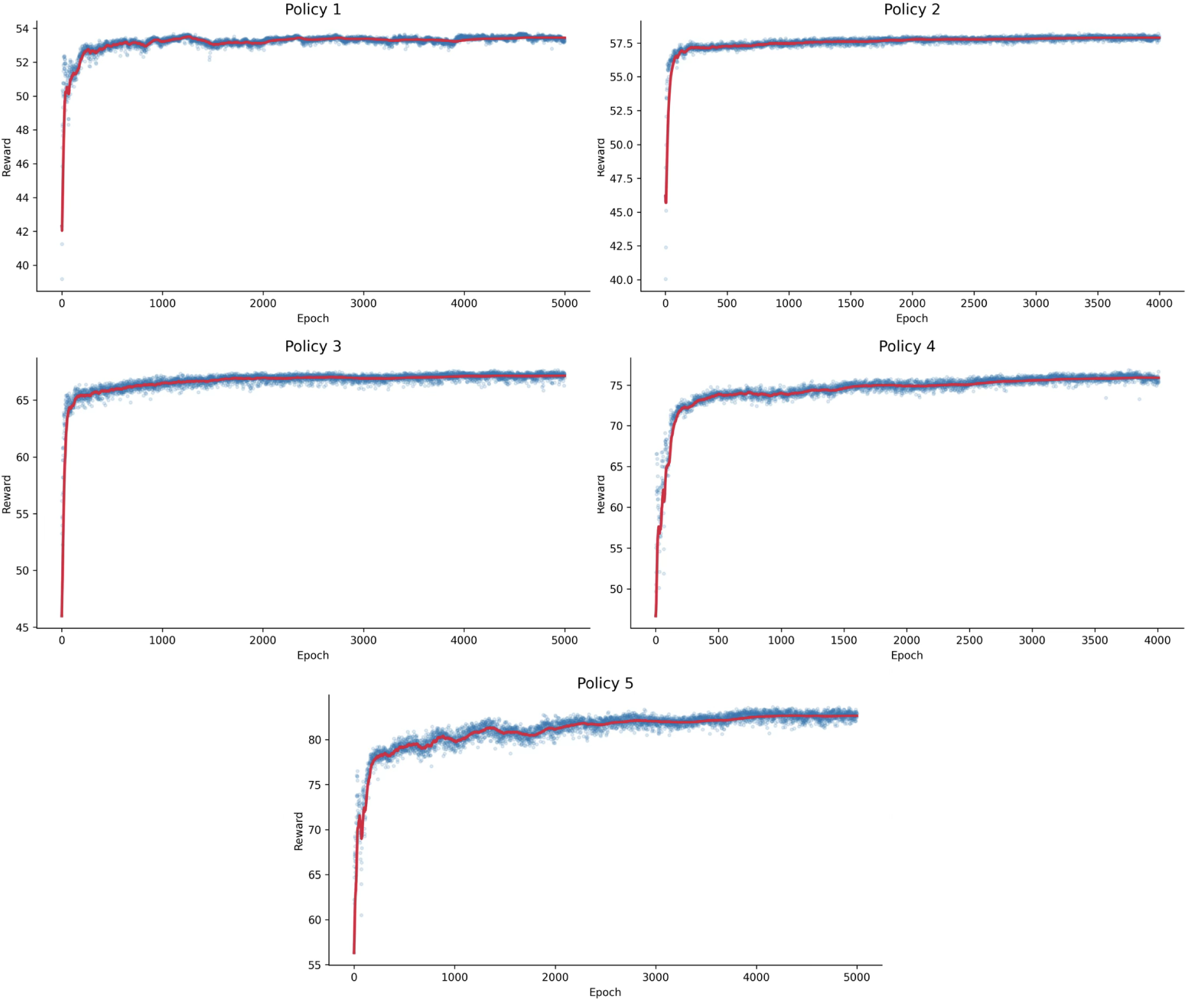
Training rewards across epochs based on a larger policy network (see *Methods*).

**Figure S6:**
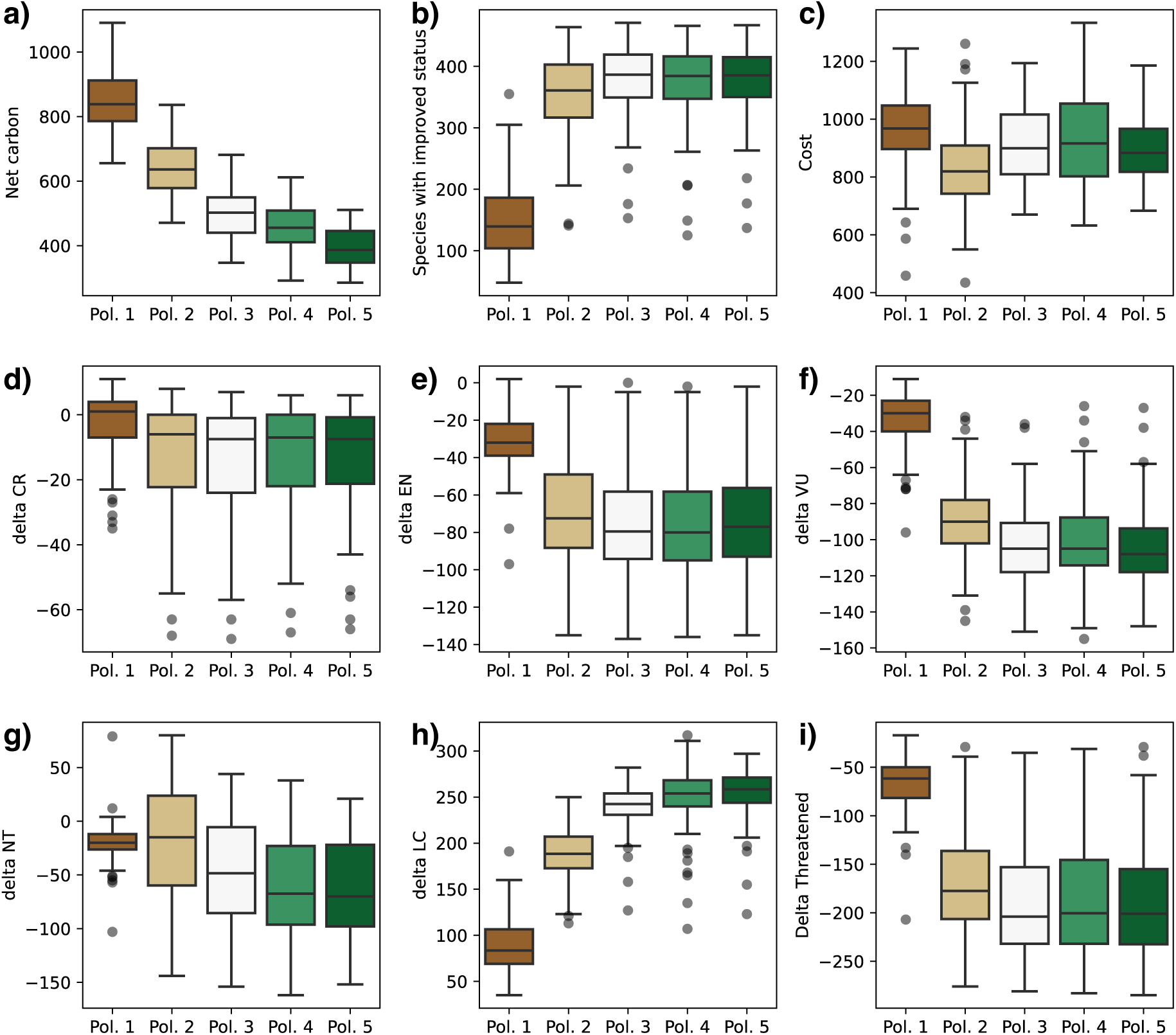
Summary statistics of key outcomes from 100 simulations run under five trained restoration models, Policies 1–5. The net carbon (a) represents the net gain in amount of carbon (here expressed in arbitrary units – see Methods) in the system after 30 time-steps during which 3,000 cells were restored. As the weight assigned to the carbon reward was lowered from 0.5 to 0 in favor of the biodiversity reward, the net carbon gain gradually decreased, although all restoration policies resulted in a net gain. In contrast, the number of species with improved conservation status, i.e. lower extinction risk, drastically increased as soon as a non-zero weight was assigned to the biodiversity reward (Policies 2–5; b). The implementation cost was similar across all policies (c), reflecting the fact that all models were trained with the same weight on the budget reward. Plots d–h) show the difference between the number of species in each extinction risk category, going from highest risk (CR) to lowest (LC), at the beginning of the simulation and the corresponding number at the end of the simulation. Thus, negative numbers, here observed in d–g) mean that the number of species in the corresponding risk categories decreased. Positive numbers, here mostly observed in h), mean that the number of species in these categories increased. These plots show that all policies that considered biodiversity rewards in the optimization (Policies 2–5) resulted in substantial reductions of threatened species compared to a policy targeting carbon only. This is also evident when lumping together all threatened species (i).

**Figure S7:**
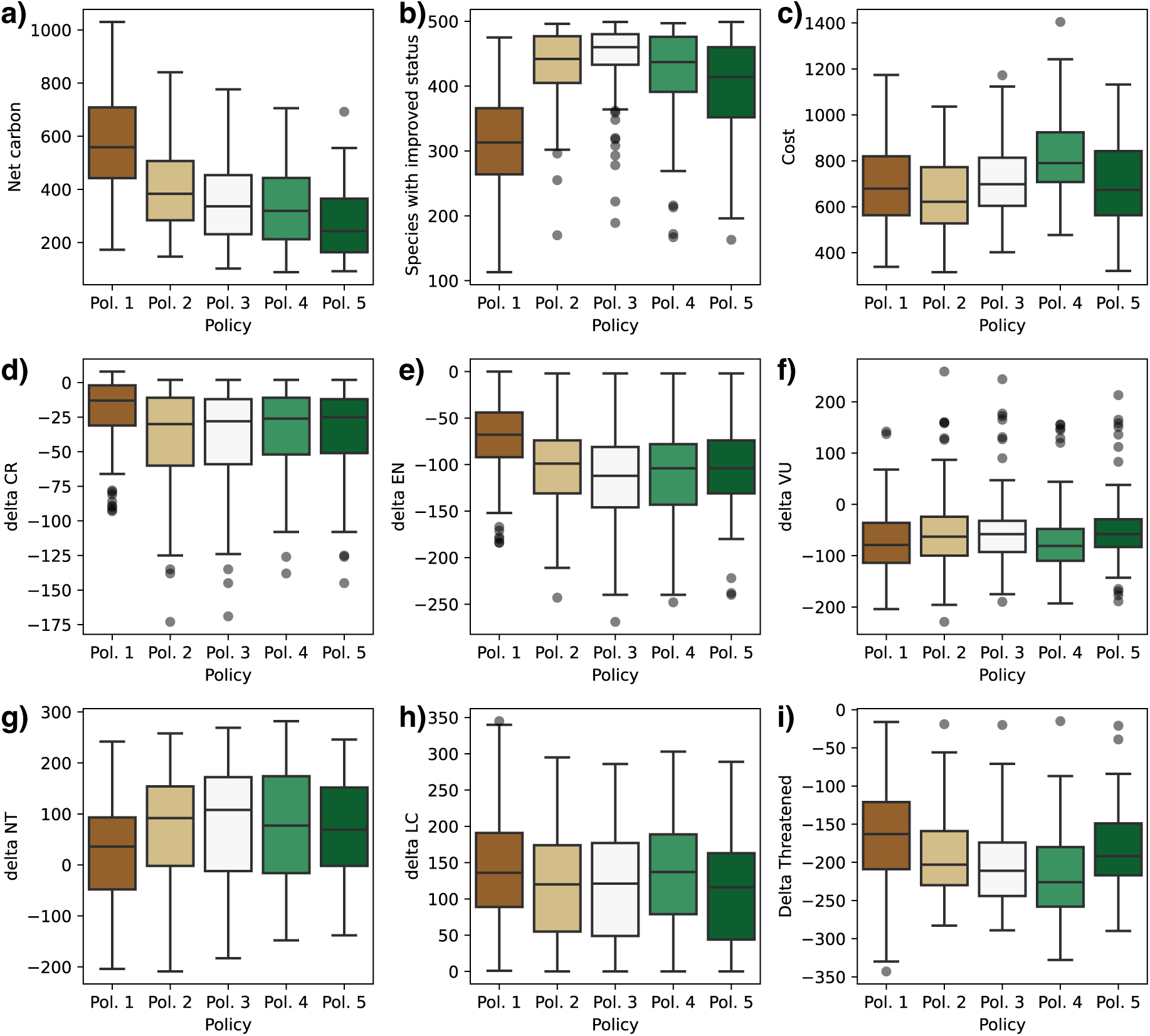
Summary of 100 simulations from random locations and variable resolution (Fig. S3) run under five trained restoration models, Policies 1–5. The general trends in terms of net carbon gain (decreasing from Policy 1 to 5; a), number of species with improved conservation status (higher from Policies 2–5; b), and costs (roughly constant across all policies) are similar to those observed across the simulations based on a fixed geographic extent, upon which the models were optimized. However the patterns are less consistent when looking at changes in species number across risk category (from CR to LC; d–h) or moving from a threatened to a non-threatened classification (i). These results suggest that the models (here trained on a single region) could be re-optimized or fine-tuned to best fit different regions.

**Figure S8:**
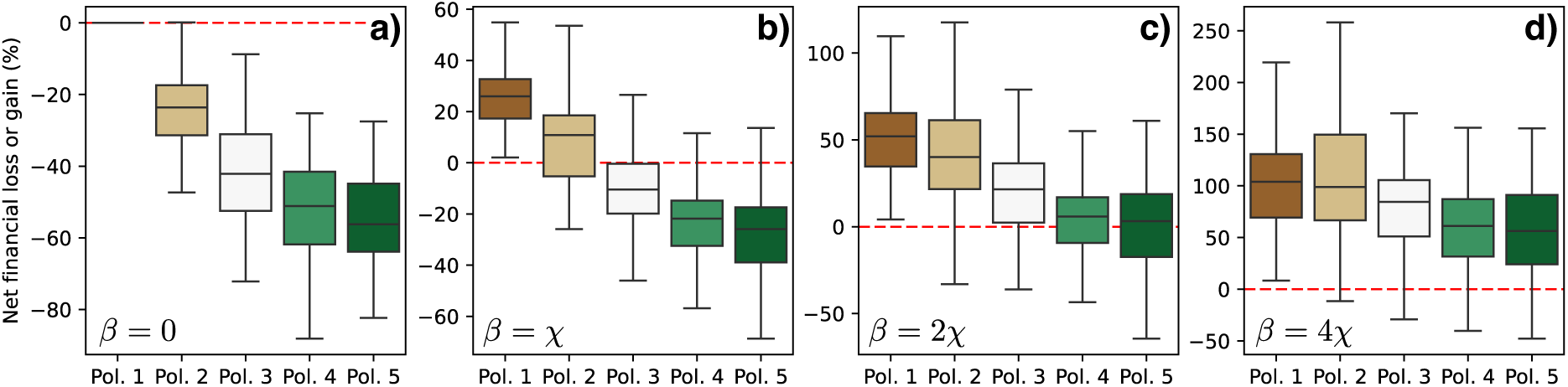
Financial outcomes of restoration policies (net percentage loss or gain, i.e. Φ in Equation 15) under different valuations of biodiversity credits, obtained from running the simulations across 100 random regions with different spatial location, extent and resolution. Negative values indicate a net economic loss where restoration costs exceed the cumulative value of carbon and biodiversity credits, while positive values indicate a net gain. The results are shown when only carbon credits (*χ*) are included (i.e. *β* = 0; a), which we used as a baseline of net 0 financial outcome to show how biodiversity credits could contribute value to the simulated restoration programs. The results in b–d) show that when biodiversity credits are valued at *β* = 4*χ*, all policies become financially viable (Φ *>* 0).

**Figure S9:**
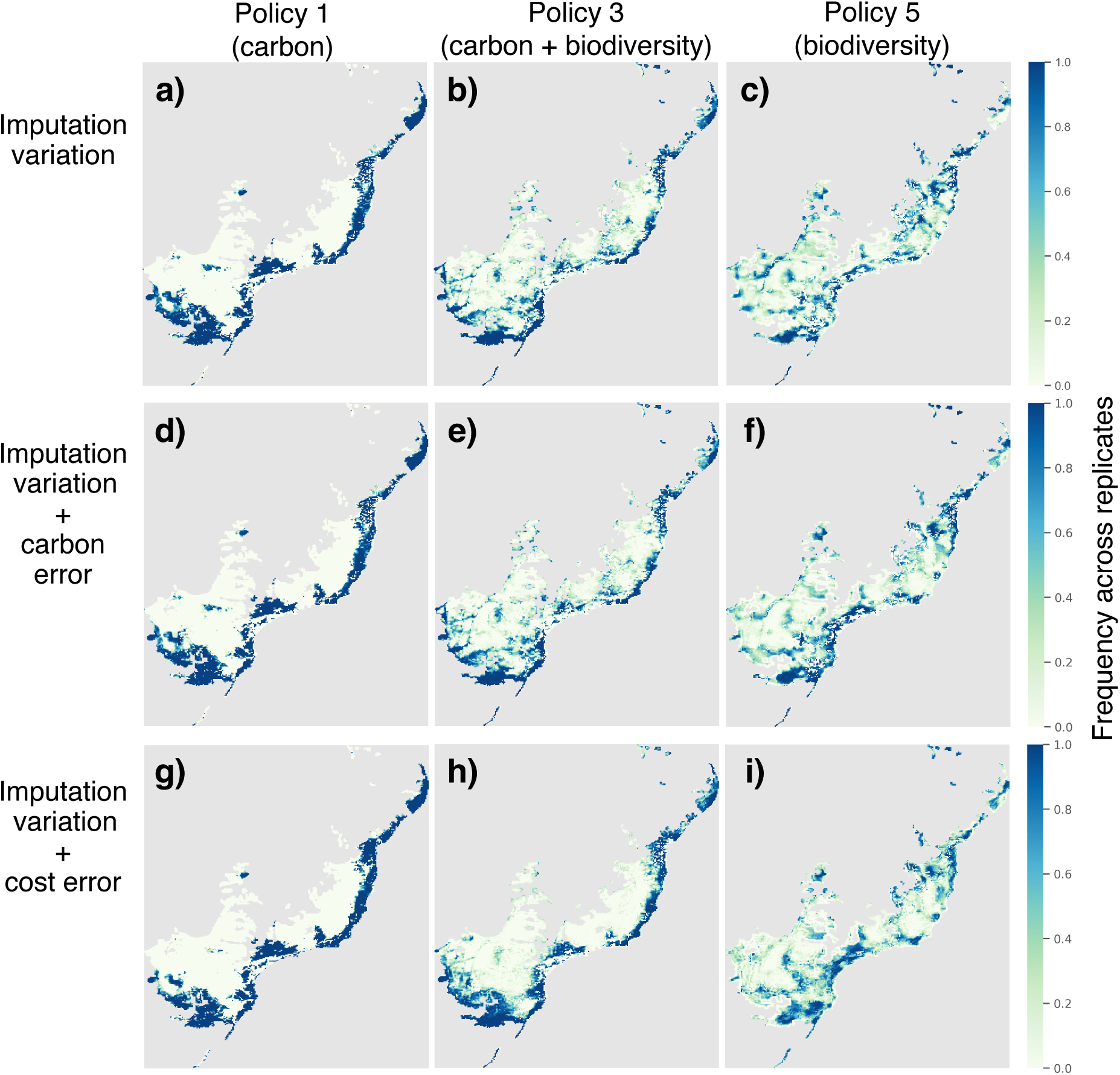
Restoration priorities in the Atlantic Forest: results for the three policies show the frequency at which each grid cell was selected for restoration (green-blue shades) across 30 replicates, where growth rates, species sensitivity to disturbance, and missing extinction risk assessments were randomly imputed (a–c). Sensitivity tests show the impact of simulated error in per-species carbon (d–f) and in the spatial distribution of restoration costs (g–i).

**Figure S10:**
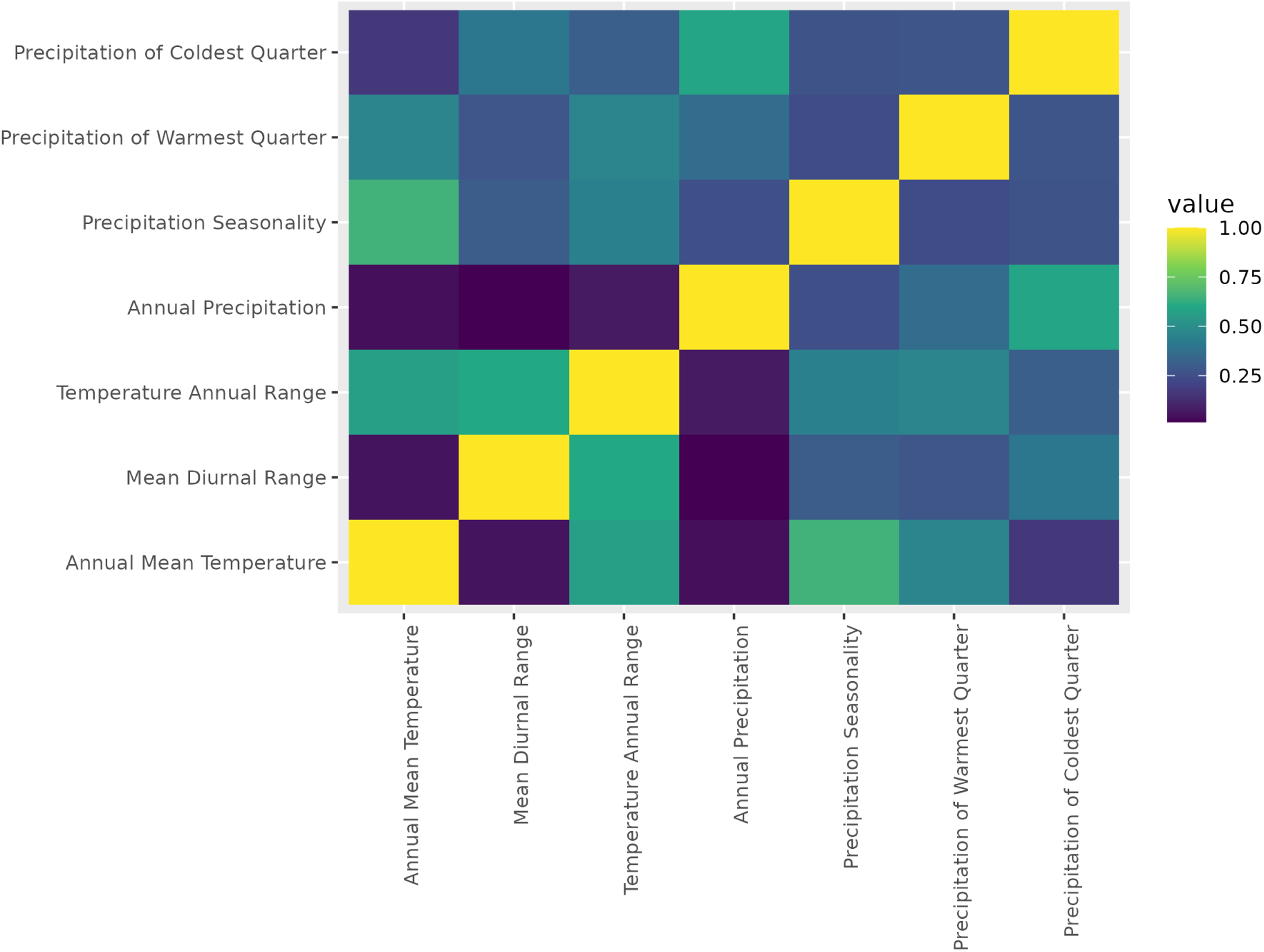
Correlation coefficients for the seven climatic variables ^69^ retained for species distribution modeling.

**Figure S11:**
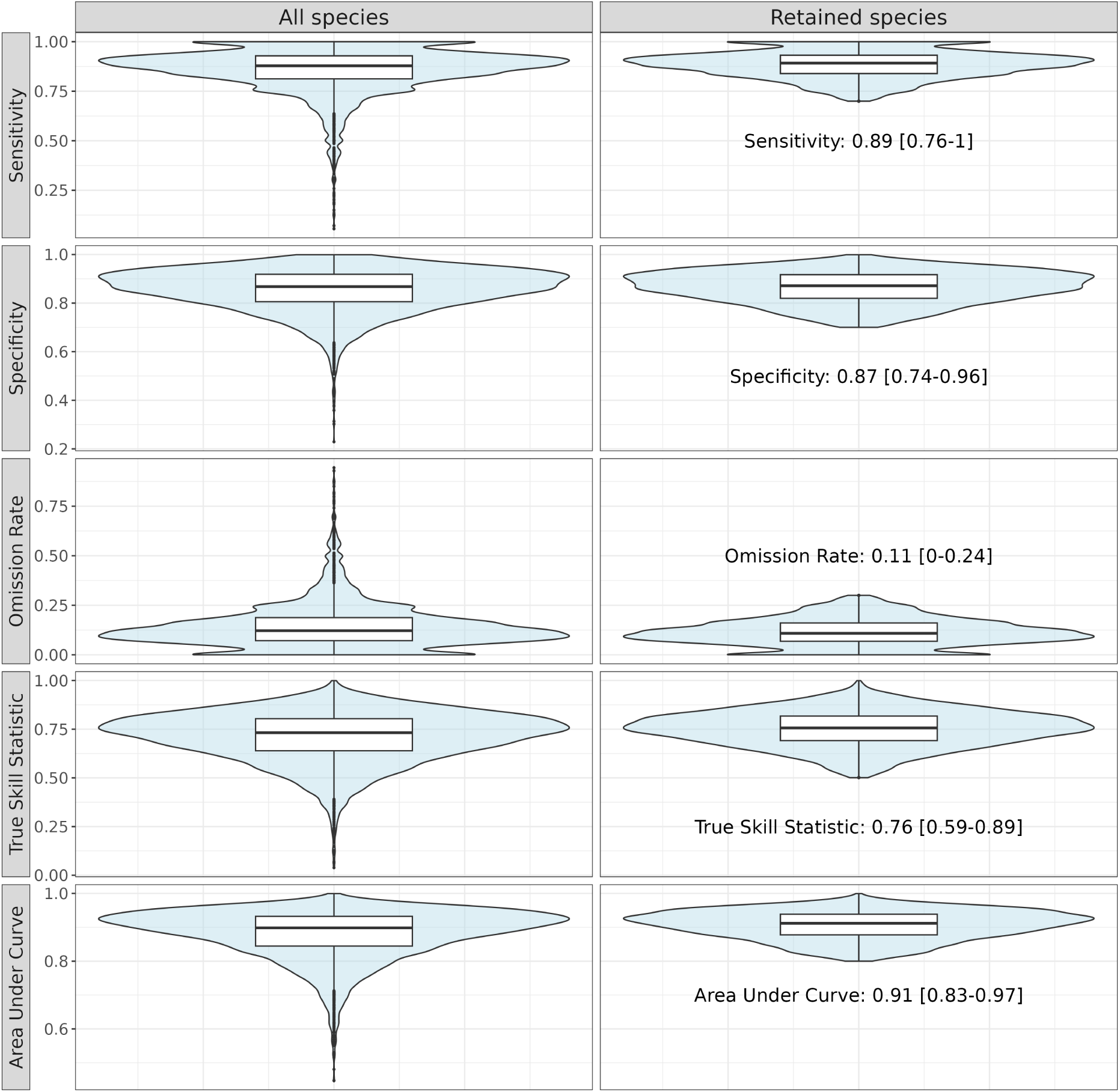
Violin plots and box plots of the five performance metrics of species distribution models across (a) all species and (b) retained species after applying exclusion criteria (sensitivity *<* 0.7, specificity *<* 0.7, True Skill Statistic *<* 0.5, and Area Under Curve *<* 0.8).

**Figure S12:**
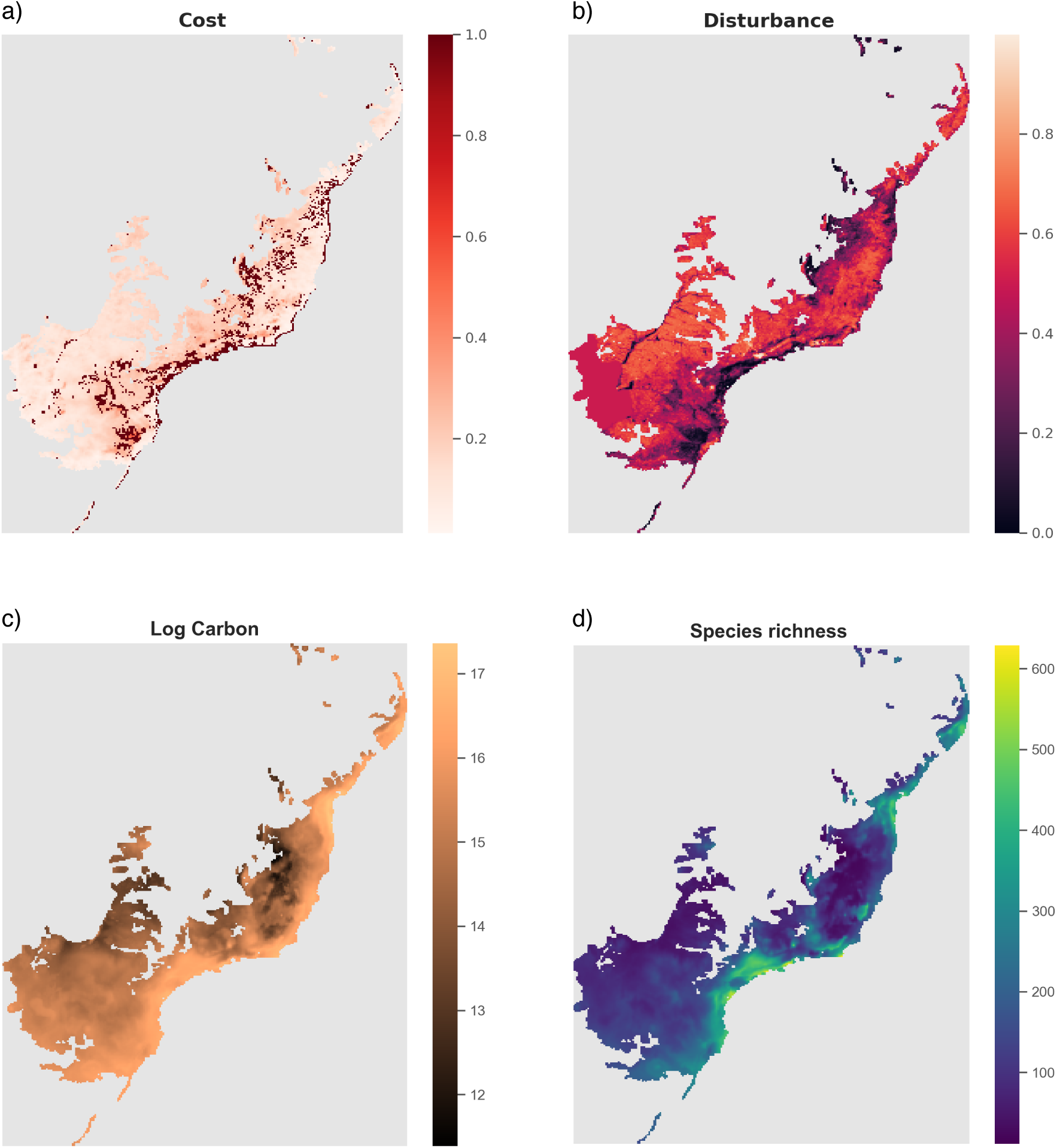
Approximate values of a) restoration costs, b) anthropogenic disturbance, c) simulated total carbon in the absence of disturbance, and d) simulated species richness based on stacking the species distribution models. The values are approximated as described in the *Methods*.

**Figure S13:**
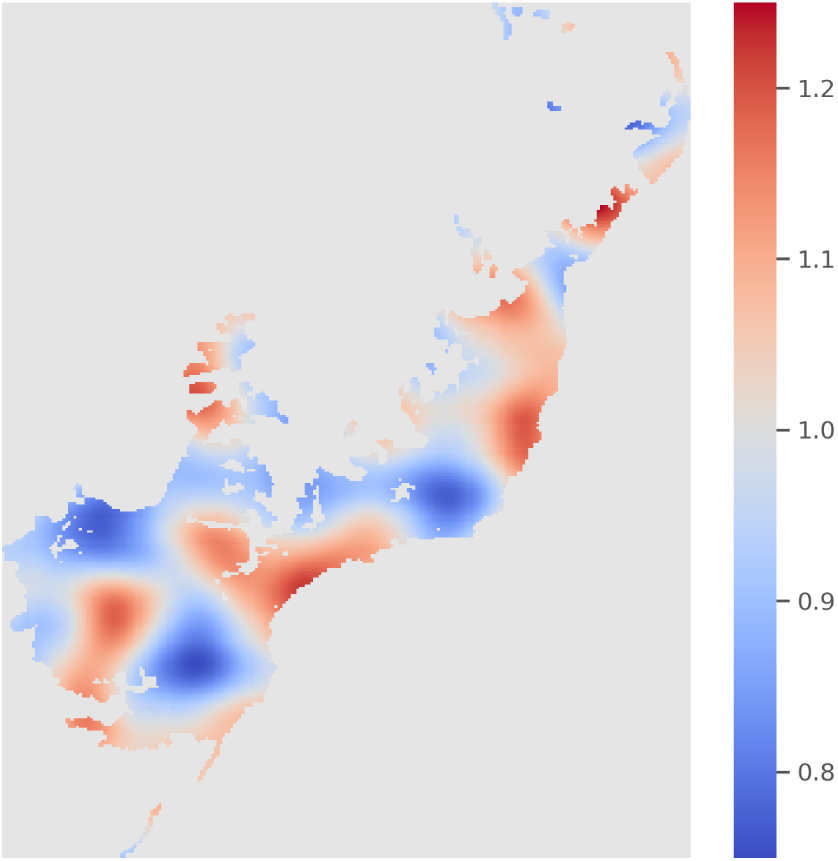
Example of Perlin noise applied as random cost multiplier to evaluate the impact of error in the cost layer on the inferred restoration priorities. The multiplier is rescaled to alter the cost layer by ± 25%.

## Supplementary Tables

**Table S1:** Biodiversity credit valuation estimated to reach the same net financial gain as in Policy 1 (focused on carbon sequestration) across all other policies (with increasing weight associated with extinction risks), computed based on Equation 17. The values are calculated across 100 simulations based on the reference region (on which models were trained).

| | $\beta/\chi$ | | | |
| --- | --- | --- | --- | --- |
|  | median | st. dev. | 25% | 75% |
| Policy 1 | 0 | 0 | 0 | 0 |
| Policy 2 | 0.80 | 2.25 | 0.51 | 1.20 |
| Policy 3 | 2.65 | 3.93 | 1.72 | 4.17 |
| Policy 4 | 2.94 | 7.66 | 2.17 | 5.01 |
| Policy 5 | 3.51 | 6.05 | 2.56 | 5.39 |

**Table S2:** Biodiversity credit valuation estimated to reach the same net financial gain as in Policy 1 across all policies, computed based on Equation 17. The values are calculated across 100 simulations based on a set of random regions differing in size and location from the region used to train the policies (Fig. S3).

| | $\beta/\chi$ | | | |
| --- | --- | --- | --- | --- |
|  | median | st. dev. | 25% | 75% |
| Policy 1 | 0 | 0 | 0 | 0 |
| Policy 2 | 2.64 | 18.09 | 0.26 | 6.50 |
| Policy 3 | 4.03 | 45.29 | 1.31 | 10.19 |
| Policy 4 | 6.06 | 13.58 | 2.09 | 11.67 |
| Policy 5 | 4.49 | 30.96 | 0.00 | 11.20 |

**Table S3:** Summary of the outcomes of 100 simulations performed under different restoration models. Non-parametric models 1–3 (see *Supplementary Methods and Results*) were based on simple metrics targeting carbon and costs (1); carbon, biodiversity and costs (2); and biodiversity and costs (3). The other models (Policies 1–5) were based on CAPTAIN-trained neural networks of 3 and 2 nodes in two hidden layers (small NN) and 16 and 8 nodes (larger NN). The effectively protected species quantify the number of species that started from a threatened category (CR, EN, VU) at the beginning of the simulations but moved to a non-threatened category (NT, LC) after restoration, while the species with improved status are species that moved to a lower extinction risk. All values are given as median among simulations and 2.5 and 97.5% quantiles rounded to integer.

|  | Model | Cost<br>median (q2.5, q97.5) | Sp. in restored areas<br>median (q2.5, q97.5) | Net carbon<br>median (q2.5, q97.5) |
| --- | --- | --- | --- | --- |
| non-parametric | 1 | 941 (591, 1260) | 383 (227, 449) | 846 (688, 1021) |
|  | 2 | 1128 (846, 1723) | 431 (339, 484) | 690 (528, 929) |
|  | 3 | 1096 (789, 1700) | 452 (364, 488) | 629 (469, 879) |
| small NN | Pol. 1 | 968 (665, 1225) | 323 (218, 437) | 838 (676, 1026) |
|  | Pol. 2 | 820 (599, 1150) | 479 (462, 489) | 636 (495, 815) |
|  | Pol. 3 | 899 (718, 1128) | 483 (475, 491) | 502 (391, 660) |
|  | Pol. 4 | 916 (720, 1278) | 482 (470, 492) | 456 (337, 597) |
|  | Pol. 5 | 883 (723, 1131) | 484 (476, 492) | 387 (307, 503) |
| larger NN | Pol. 1 | 1015 (693, 1316) | 406 (235, 467) | 809 (646, 1033) |
|  | Pol. 2 | 817 (654, 1065) | 476 (457, 488) | 653 (513, 918) |
|  | Pol. 3 | 833 (602, 1064) | 483 (469, 491) | 532 (404, 798) |
|  | Pol. 4 | 885 (658, 1205) | 481 (468, 491) | 500 (382, 718) |
|  | Pol. 5 | 924 (705, 1178) | 483 (474, 492) | 448 (340, 628) |
|  | Model | Effectively protected sp.<br>median (q2.5, q97.5) | Sp. with improved status<br>median (q2.5, q97.5) |  |
| non-parametric | 1 | 73 (32, 159) | 172 (85, 317) |  |
|  | 2 | 119 (48, 231) | 263 (119, 415) |  |
|  | 3 | 133 (52, 233) | 295 (131, 435) |  |
| small NN | Pol. 1 | 62 (23, 125) | 140 (62, 272) |  |
|  | Pol. 2 | 178 (63, 247) | 361 (218, 453) |  |
|  | Pol. 3 | 204 (71, 268) | 387 (250, 463) |  |
|  | Pol. 4 | 201 (56, 274) | 385 (206, 458) |  |
|  | Pol. 5 | 201 (65, 268) | 386 (239, 463) |  |
| larger NN | Pol. 1 | 84 (51, 145) | 207 (136, 286) |  |
|  | Pol. 2 | 165 (55, 237) | 352 (197, 441) |  |
|  | Pol. 3 | 201 (61, 259) | 383 (237, 454) |  |
|  | Pol. 4 | 205 (65, 282) | 363 (203, 444) |  |
|  | Pol. 5 | 208 (61, 279) | 385 (235, 455) |  |

